# Genome-resolved viral and cellular metagenomes revealed potential key virus-host interactions in a deep freshwater lake

**DOI:** 10.1101/655167

**Authors:** Yusuke Okazaki, Yosuke Nishimura, Takashi Yoshida, Hiroyuki Ogata, Shin-ichi Nakano

## Abstract

Metagenomics has dramatically expanded the known virosphere, but freshwater viral diversity and their ecological interaction with hosts remain poorly understood. Here, we conducted a metagenomic exploration of planktonic dsDNA prokaryotic viruses by sequencing both virion (<0.22 μm) and cellular (0.22–5.0 μm) fractions collected spatiotemporally from a deep freshwater lake (Lake Biwa, Japan). This simultaneously reconstructed 183 complete (i.e., circular) viral genomes and 57 bacterioplankton metagenome-assembled genomes. Analysis of metagenomic read coverage revealed vertical partitioning of the viral community analogous to the vertically stratified bacterioplankton community. The hypolimnetic community was generally stable during stratification, but occasionally shifted abruptly, presumably due to lysogenic induction. Genes involved in assimilatory sulfate reduction were encoded in 20 (10.9%) viral genomes, including those of dominant viruses, and may aid viral propagation in sulfur-limited freshwater systems. Hosts were predicted for 40 (21.9%) viral genomes, encompassing 10 phyla (or classes of Proteobacteria) including ubiquitous freshwater bacterioplankton lineages (e.g., *Ca.* Fonsibacter and *Ca.* Nitrosoarchaeum). Comparison with viral genomes derived from published metagenomes revealed viral phylogeographic connectivity in geographically isolated habitats. Notably, analogous to their hosts, actinobacterial viruses were among the most diverse, ubiquitous, and abundant viral groups in freshwater systems, with potential high lytic activity in surface waters.

## Introduction

Since the discovery three decades ago that a large number of viruses are present in aquatic systems (Bergh *et al.*, 1989), our understanding of viral ecology has grown in an unprecedented manner. Whereas viruses can infect cells of all three domains of life, those infecting bacteria and archaea have been studied extensively, as they are numerically dominant and play important roles in their ecosystems. Such viruses rewire the microbial food web by releasing the host’s cellular contents back into the environment (the viral shunt) (Wilhelm and Suttle, 1999) and may carry host-derived metabolic genes (auxiliary metabolic genes; AMGs), which amend the host’s metabolism to facilitate viral propagation (Breitbart *et al.*, 2007). They are key players in maintaining the diversity of the microbial community by selectively killing the dominant hosts (Thingstad, 2000) or promoting the genetic diversification of hosts (Koskella and Brockhurst, 2014; Touchon *et al.*, 2017).

Although freshwater ecosystems occupy a small portion of the earth’s surface, these areas cycle petagrams of carbon per year globally (Tranvik *et al.*, 2009; Raymond *et al.*, 2013), and are of great importance to humans as drinking water sources. Furthermore, freshwater ecosystems are vulnerable to climate change (Posch *et al.*, 2012; Woolway and Merchant, 2019) and eutrophication (Jenny *et al.*, 2016). Microbial processes are key to understanding these ecological and biogeochemical mechanisms. Researchers have reached the consensus that surface freshwater systems (epilimnion) are ubiquitously dominated by bacterioplankton lineages represented by the phyla Actinobacteria, Proteobacteria, and Bacteroidetes (Newton *et al.*, 2011). In addition, we have previously unveiled the vertical stratification of bacterioplankton communities in deep freshwater lakes, where the oxygenated hypolimnion— i.e., the deep aerobic water layer below the thermocline, which generally constitutes a volumetrically significant part of deep temperate, oligo-mesotrophic lakes— was dominated by specific lineages that are uncommon in the epilimnion, including members of Chloroflexi (Okazaki *et al.*, 2013, 2018), Planctomycetes, Nitrospira and Thaumarchaeota (Okazaki and Nakano, 2016; Okazaki *et al.*, 2017). These lineages are likely responsible for important biogeochemical processes in the oxygenated hypolimnion, such as remineralization or conversion of semi-labile organic matter (Maki *et al.*, 2010; Thottathil *et al.*, 2013), nitrification (Small *et al.*, 2013; Alfreider *et al.*, 2018), and methane oxidation (Murase and Sugimoto, 2005).

The recent introduction of viral metagenomics (viromics) opened the door to elucidating the vast diversity of uncultured viral genomes and genes (Brum and Sullivan, 2015) and facilitated identification of viruses infecting predominant bacterial hosts. Through both cultivation-dependent and -independent manners, viruses that infect major freshwater bacterioplankton lineages have been documented in the last few years, including those of Actinobacteria (Ghai *et al.*, 2017), *Polynucleobacter* (Cabello-Yeves *et al.*, 2017), LD28 (*Ca.* Methylopumilus) (Moon *et al.*, 2017) and Comamonadaceae (Moon *et al.*, 2018). Elucidating such predominant virus-host interactions is key to characterizing viruses playing central roles in the microbial ecosystem and biogeochemical cycling. However, despite a growing number of viromic studies in freshwater systems (Skvortsov *et al.*, 2016; Moon *et al.*, 2017; Silva *et al.*, 2017; Arkhipova *et al.*, 2018), viruses infecting other typical freshwater bacterioplankton lineages, such as LD12 (*Ca.* Fonsibacter) (Henson *et al.*, 2018) and *Limnohabitans* (Kasalický *et al.*, 2013) have yet to be characterized. Moreover, no viromic study has investigated an oxygenated hypolimnion, leaving hypolimnetic viral diversity and virus-host interactions totally unexplored.

The main aim of this study was to characterize key virus-host interactions in freshwater ecosystems. To this end, we performed spatiotemporal (from two water layers for nine months) sampling of both the cellular (size = 0.22–5.0 μm) and virion (<0.22 μm) fractions at a pelagic station of a deep freshwater lake. This allowed simultaneous reconstruction of both viral and host (i.e., prokaryotic) genomes from the water column and to follow the lifecycles of individual viral lineages including both intra- and extra-cellular stages. The assembled genomes were compared with viral genomes derived from published metagenomic datasets, and the result demonstrated phylogeographic connectivity of the viral community among habitats. Overall, the present study provided an overview of genome-resolved viral diversity in freshwater systems and revealed virus-host interactions of potential ecological importance, including those involving ubiquitous freshwater bacterial lineages.

## Results and Discussion

### Overview of samples and assemblies

In total, 12 water samples were collected at a pelagic site on Lake Biwa, the largest freshwater lake in Japan. The epilimnion (5 m) was sampled over 3 months and the hypolimnion (65 m) was sampled over 9 months. Water temperature, dissolved oxygen concentration, and the abundances of bacteria and virus-like particles in the study site (Table S1) were within the range reported from previous studies in the lake (Pradeep Ram *et al.*, 2010; Okazaki *et al.*, 2013). Notably, thermal stratification was observed from June to December and the hypolimnion was always oxygenated, with dissolved oxygen concentrations of >6.0 mg L^−1^. Metagenomic sequencing with subsequent assembly (Table S2), decontamination, and dereplication steps eventually yielded 4,158 viral contigs of >10 kb (Lake Biwa Viral Contigs [LBVCs]), including 183 complete (i.e., circular) genomes (Lake Biwa Viral Genomes [LBVGs]) (Fig. S1). The GC content of the 183 LBVGs ranged from 29.1 to 66.7%, and all genome sizes were <120 kb except for LBVG_1 (318 kb) and LBVG_2 (190 kb) (Fig. 1A), which were within the range of known prokaryotic dsDNA viral genomes (Mahmoudabadi and Phillips, 2018).

**Figure 1.**
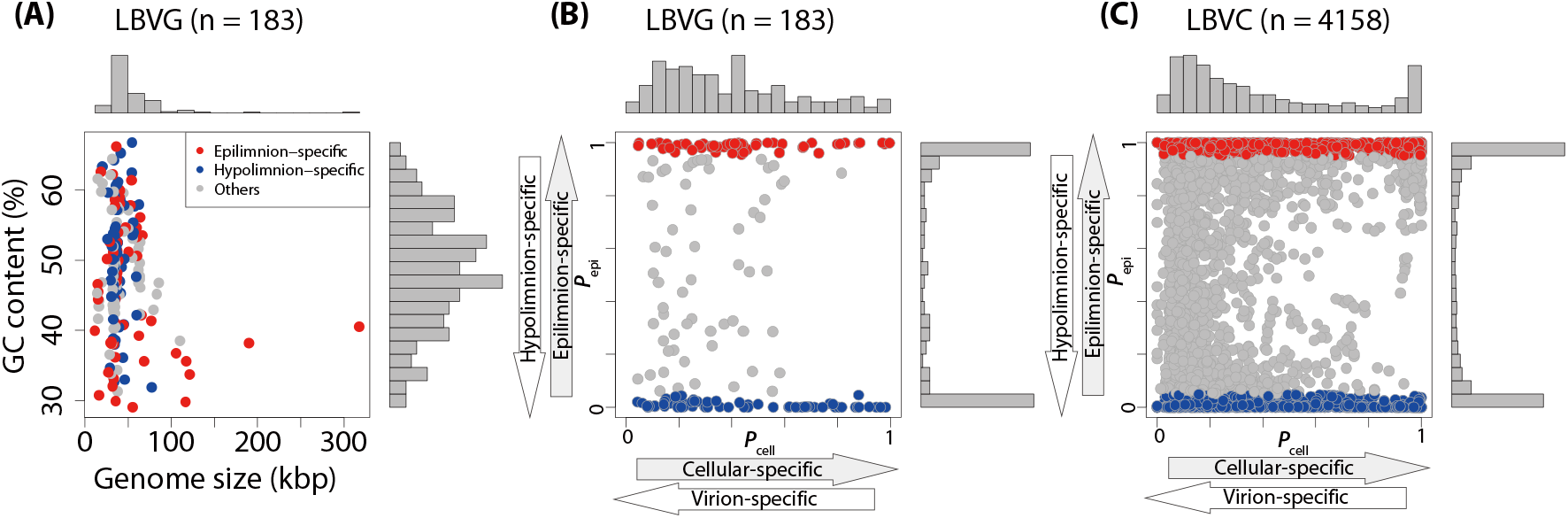
Profiles of assembled viral genomes and contigs. (A) Distribution of genome size and GC content (%) of Lake Biwa Viral Genomes (LBVGs). Habitat preferences of (B) LBVGs and (C) Lake Biwa Viral Contigs (LBVCs). The vertical and horizontal axes indicate the average *P*_epi_ and *P*_cell_ for each LBVG, respectively. Viruses with average *P*_epi_ >95% and <5% were designated as epilimnion- and hypolimnion-specific viruses and are indicated with red and blue points, respectively.

Similarity between viral genomes was evaluated using *S*_G_, a similarity measure (ranging from 0 to 1) based on genome-wide tBLASTx scores (Nishimura *et al.*, 2017a). A previous study demonstrated that *S*_G_ >0.15 was the best threshold for genus-level clustering of viral genomes (Nishimura *et al.*, 2017a). Using this threshold, the 183 LBVGs were grouped into 28 genomic operational taxonomic units (gOTUs) containing 2–10 LBVGs each, leaving 90 LBVGs ungrouped (Fig. 2). To assess the novelty or ubiquity of LBVGs, published viral genomic sequences were compiled into two databases— the Reference Viral Genome (RVG) and Environmental Viral Genome (EVG). The RVG included 2,621 isolated prokaryotic dsDNA viral genomes. The EVG included 2,860 metagenomically recovered viral genomes and manually collected viral genomes that had been reported to infect the dominant freshwater bacterioplankton lineages, including both isolated and metagenome-assembled genomes (Fig. S2). Using the threshold of *S*_G_ >0.15, 4 (2.2%) and 127 (69.4%) LBVGs were found to have genus-level relatives in the RVG and EVG, respectively (Fig. 2). An *S*_G_- based proteomic tree including all LBVGs and EVGs and alignments among genomes is available at https://www.genome.jp/viptree/u/LBV/retree/LBVGandEVG.

**Figure 2.**
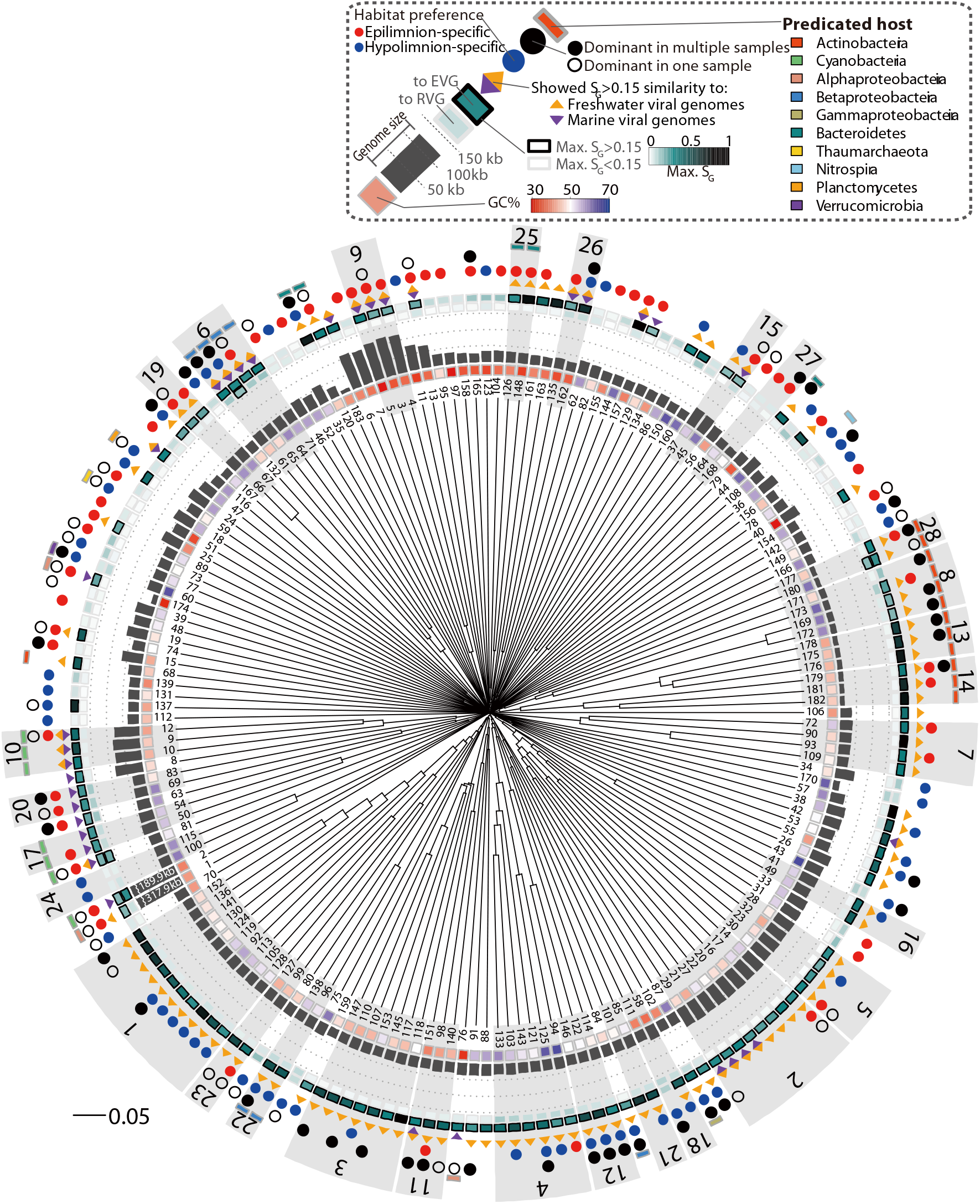
Proteomic tree of 183 LBVGs with summarized analytical results. The gray shaded boxes indicate individual gOTUs with corresponding gOTU numbers. Annotation of epilimnion- and hypolimnion-specific viruses corresponds with the definition in Figure 1. “Dominant” viruses shown by the black and white circles indicate “ranked among the 10 most abundant viruses,” as described in Figure 4.

Metagenomic binning of the contigs assembled from the cellular fraction recovered 57 metagenome-assembled genomes of bacterioplankton (Lake Biwa metagenome-assembled genomes [LBMAGs]), including diverse bacterial and archaeal phyla (Figs. 3 and S3). Forty-five (78.9%) of these genomes were of high quality, with estimated completeness and contamination scores of >80% and <5%, respectively (Table S3). Consistent with previous observations (Okazaki and Nakano, 2016; Okazaki *et al.*, 2017), a depth-stratified bacterioplankton community was observed— members of LD12, Actinobacteria, and Cyanobacteria were dominant in the epilimnion, whereas CL500-11 and *Ca.* Nitrosoarchaeum were dominant in the hypolimnion (Table S4). In several lineages (e.g., LD12, LD28, and OPB56), distinct genomes were recovered from the epilimnion and hypolimnion, indicating micro-diversification between the water layers. Two LBMAGs representing candidate phyla, *Ca*. Levybacteria (OP11) and *Ca*. Saccharibacteria (TM7) (Brown *et al.*, 2015), had not been reported in a previous study conducted at Lake Biwa using 16S rRNA gene amplicon sequencing (Okazaki and Nakano, 2016), as these phyla harbor 16S rRNA gene sequences that are undetectable with conventional universal primers.

**Figure 3.**
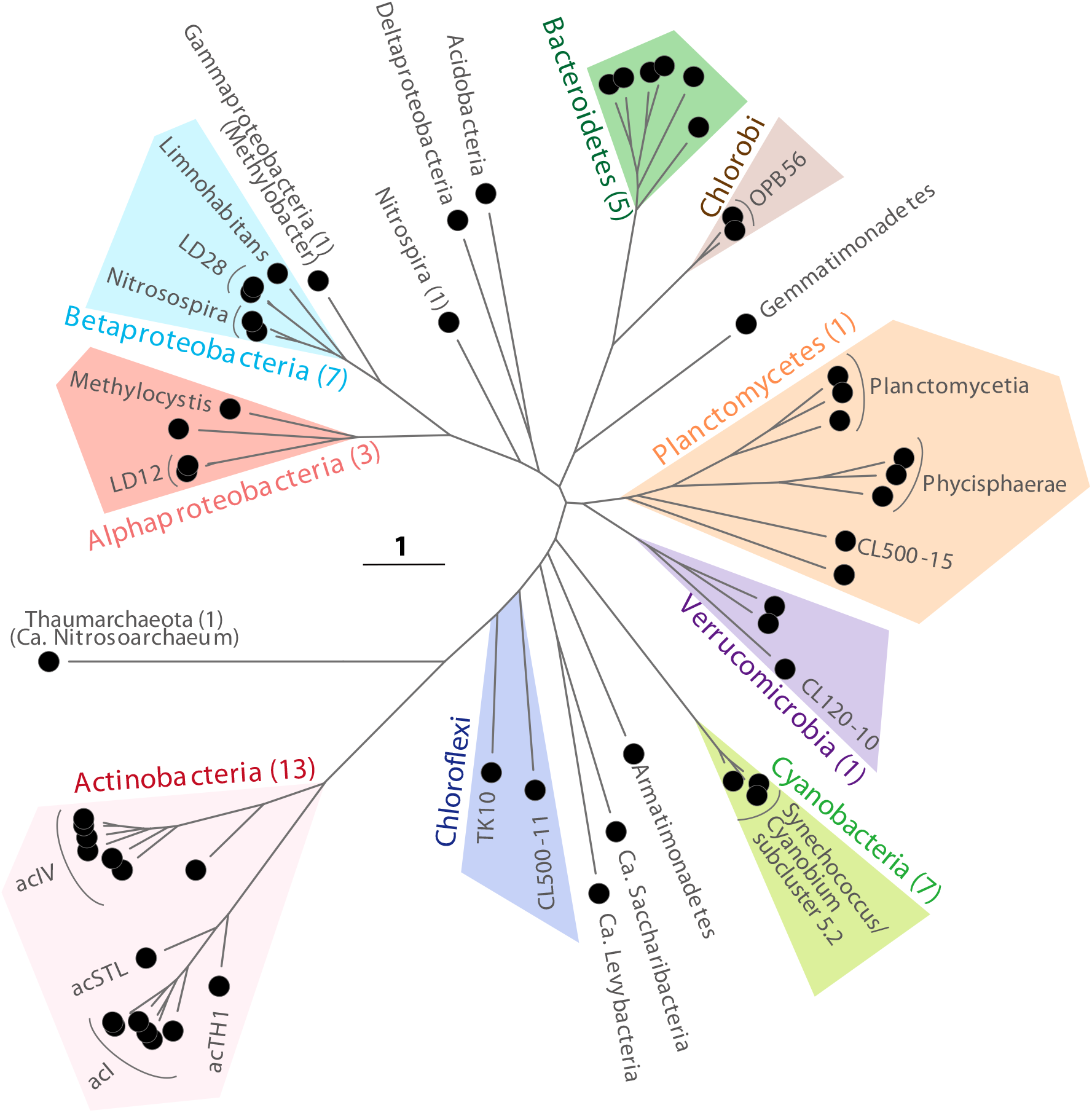
A genome-resolved phylogenetic tree of Lake Biwa metagenome-assembled genomes (LBMAGs) based on conserved single-copy genes selected using PhyloPhlAn software. Members of the same phylum (or Proteobacteria class) were grouped by the same color shade. Numbers in parentheses indicate the number of LBVGs predicted to infect each taxonomic group. Note that Chloroflexi did not form a monophyletic clade here but did in a tree that included reference genomes (Fig. S3).

### Spatiotemporal dynamics of the viral community

Relative abundance of LBVGs and LBVCs was determined based on metagenomic read coverage, shown as fragments per kilobase per mapped million reads (FPKM) (Nishimura *et al.*, 2017a; Yoshida *et al.*, 2018). The result revealed that the epilimnion- and hypolimnion-specific viral lineages were clearly separated (Fig. 1B, C). Among the 183 LBVGs, 60 were epilimnion-specific and 62 were hypolimnion-specific, demonstrating epilimnion-hypolimnion preference (*P*_epi_) values of >95% and <5%, respectively. Non-metric multidimensional scaling (NMDS) analysis further demonstrated vertical separation of the viral communities during the stratification period (Fig. 4). The depth-stratified viral community is analogous to that reported from ocean (Hurwitz *et al.*, 2015; Mizuno *et al.*, 2016; Paez-Espino *et al.*, 2016; López-Pérez *et al.*, 2017; Luo *et al.*, 2017) and likely reflects the depth-stratified community of bacterioplankton (Okazaki and Nakano, 2016) (Table S4).

**Figure 4.**
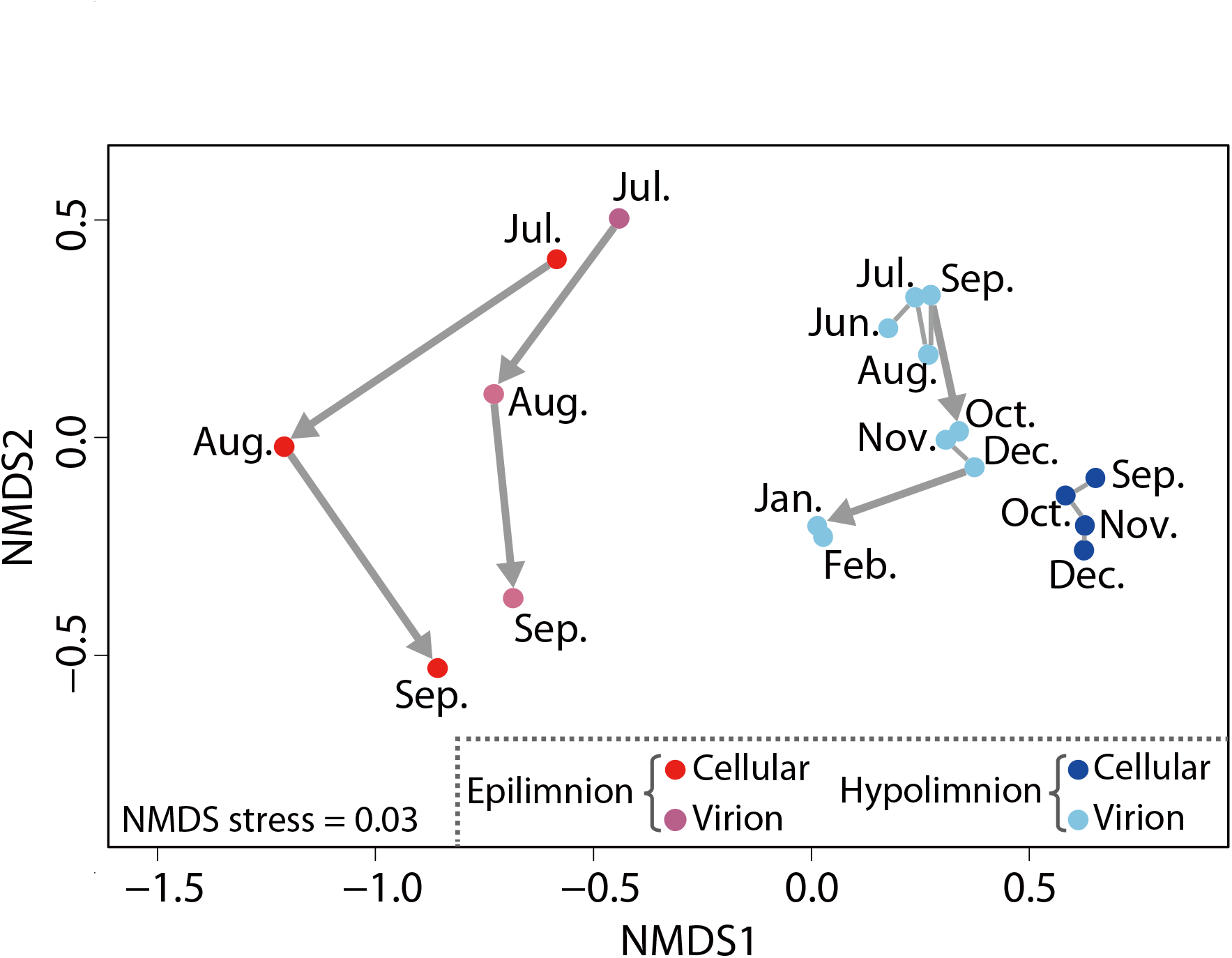
Non-metric multidimensional scaling (NMDS) analysis showing dissimilarity of the viral community between samples. The distance between samples was calculated as Bray-Curtis dissimilarity based on the composition of the 183 LBVGs. Gray lines connect samples of consecutive months in the same sample fraction, where bold arrows indicate a large community shift with the Bray-Curtis dissimilarity of >0.5.

The viral community was generally stable in the hypolimnion (the Bray-Curtis dissimilarity of the communities between consecutive months = 0.21–0.34; except for September to October and December to January as discussed later) compared to the epilimnetic community (0.54–0.68) (Figs. 4 and 5). The rapid succession of the viral community in the epilimnion was in line with previous studies (Rodriguez-Brito *et al.*, 2010; Arkhipova *et al.*, 2018). By contrast, in the hypolimnion, many viral lineages persistently dominated for more than a month (Fig. 5). Notably, the decay rate of virioplankton ranges from 0.14 to 54% h^−1^ (half-life = 0.9 h to 20.6 days) (Wommack and Colwell, 2000), meaning that most virions in the water column turn over within a month. In addition, a recent study in a marine system reported that the majority of metagenomically-detected viruses were transcriptionally active within a period of 24 h (Yoshida *et al.*, 2018). Consequently, the continued dominance of some viral lineages in the hypolimnion likely resulted from the continuous production of virions, rather than their carryover for more than a month. This pattern might be attributable to the lower production of bacterioplankton in the hypolimnion, resulting in a prolonged viral lytic cycle. To support this, a previous study in Lake Biwa estimated that bacterial production was 10-fold lower in the hypolimnion (0.4 ± 0.1 µg C l^−1^ d^−1^) than in the epilimnion (4.2 ± 3.2 µg C l^−1^ d^−1^), and the percentage of daily bacterial production lost to viruses was 4-fold lower in the hypolimnion (13.6 ± 5.2%) than in the epilimnion (52.7 ± 16.2%) (Pradeep Ram *et al.*, 2010).

**Figure 5.**
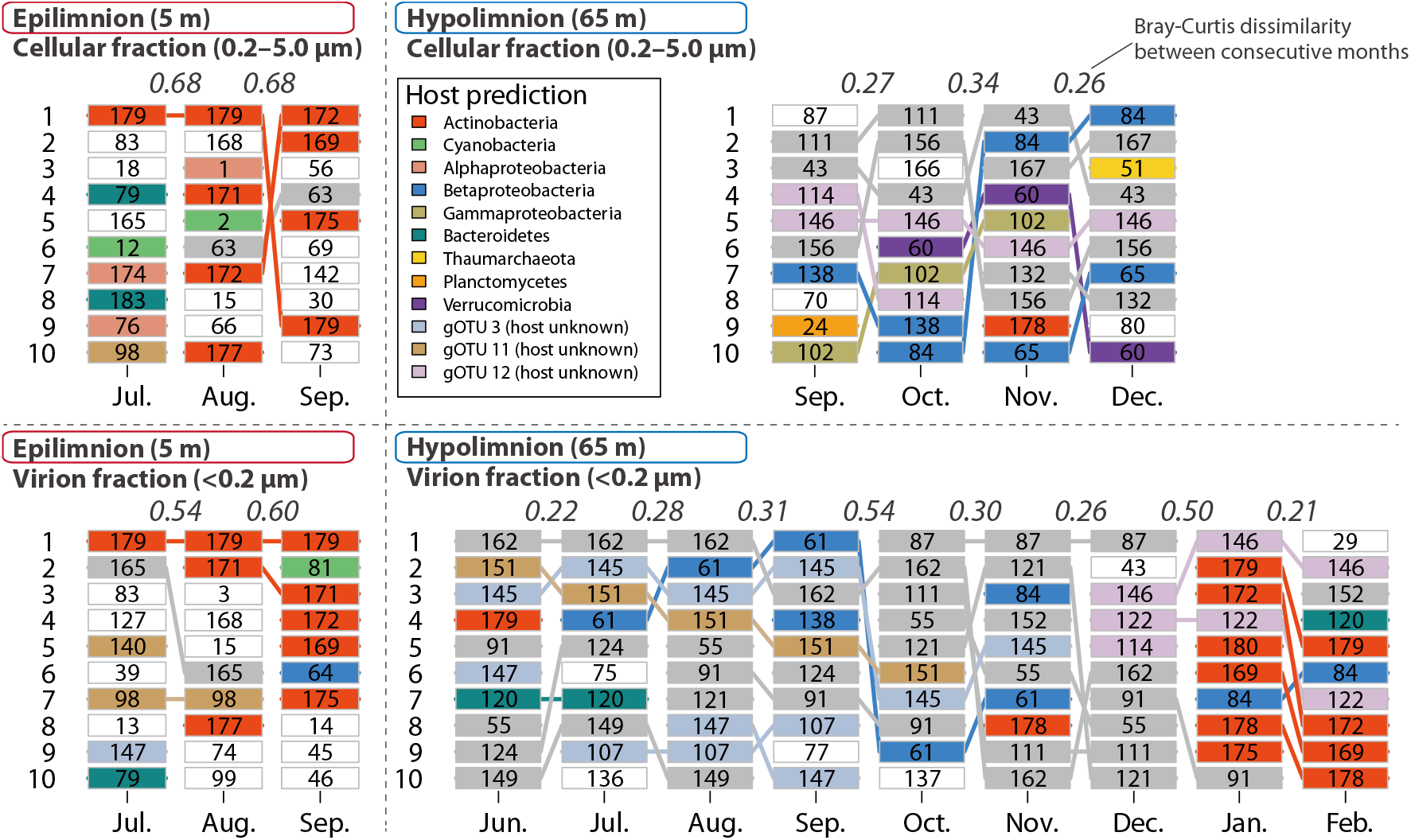
Succession of the 10 most abundant viral genomes in each sample fraction. The number in each box corresponds to the number of LBVGs. The color of each box indicates the phylum of the predicted host. The abundant gOTUs (gOTU_3, gOTU_11, and gOTU_12) are also shaded, although their hosts remain unknown. Otherwise, members that were dominant over multiple months are in gray boxes and those that were dominant only within one month are in white boxes. Italicized numbers at the top of each panel indicate the extent of the community composition shift in consecutive months, based on the Bray-Curtis dissimilarity of the whole community. Quantitative data for this figure are available in Figure S4.

In addition to the overall stability of the viral community in the hypolimnion, two dynamic community shifts were observed in the virion fraction, from September to October (Bray-Curtis dissimilarity = 0.54) and from December to January (0.50) (Figs. 4 and 5). The first shift could be resulted from several viruses switching from lysogeny to lysis (i.e., induction). In the cellular fraction, LBVG_87 was the most abundant LBVG in September but its abundance decreased abruptly in October, whereas its abundance in the virion fraction increased 120-fold over the same period, making LBVG_87 the most abundant virioplankton (Figs. 5, S4 and S5). Similarly, the relative abundances of LBVG_111 and LBVG_121 in the virion fraction increased >1800-fold and 38-fold from September to October, respectively, likely due to the induction of abundant intracellular viruses in September (Figs. 5, S4 and S5). Regrettably, the host for LBVG_87, LBVG_111, and LBVG_121 could not be identified. Since no integrase gene was found in their genomes, they may have existed in a host’s cell in an extrachromosomal or pseudolysogenic manner. As proposed for the deep ocean (Weinbauer *et al.*, 2003; Paul, 2008; Mizuno *et al.*, 2016; Luo *et al.*, 2017), the lysogenic strategy in deeper water likely results from low bacterioplankton productivity, which is insufficient to support lytic viral propagation. Lytic induction can be triggered by environmental stressors such as pH, temperature, nutrients, oxidative stress, and solar radiation (Weinbauer, 2004; Howard-Varona *et al.*, 2017). While the hypolimnion during stratification period is more physicochemically stable than the epilimnion, biological factors, e.g., a pulse of sinking phytoplankton cells from the epilimnion (Kagami *et al.*, 2006), could have resulted in a monthly-scale environmental shift and triggered the induction in the hypolimnion. Induction can also be trigged by the increasing cell abundance or improved physiological condition of the host, with the virus acting as a “time bomb” and breaking the symbiotic relationship with their host once the host has successfully bloomed in the environment (Paul, 2008; Brum *et al.*, 2016). Indeed, hypolimnion-specific bacterial lineages often take months to archive their peak abundance (Okazaki *et al.*, 2013, 2018; Okazaki and Nakano, 2016), suggesting that lysogeny (or presudolysogeny) is preferred strategy for viruses during the long growth phase of the host. No difference in cellular-virion preference (*P*_cell_) was observed between water layers (Fig. S6). Thus, lysogeny and induction may also be a common strategy in the epilimnion, although temporal trends were not clearly observed in that layer, presumably due to the rapid succession of the epilimnetic virion community (Figs. 4 and 5), which could not be fully resolved through monthly monitoring.

The second shift in the virion community from December to January likely resulted from the onset of winter vertical mixing (Table S1), which intermingled the epilimnetic and hypolimnetic viral communities, as observed for bacterioplankton (Okazaki *et al.*, 2013, 2018; Okazaki and Nakano, 2016). Alternatively, environmental stresses associated with the mixing event (e.g., increased solar radiation) could have induced lysogens in the lytic cycle (Winter *et al.*, 2018) and resulted in the observed viral community shift.

### Possible metabolic reprogramming of hosts by viral AMGs

Functional annotation of viral genes (Supplementary Dataset S1) revealed that LBVGs encoded AMGs that have previously been identified in marine viromes (Hurwitz *et al.*, 2015; Hurwitz and U’Ren, 2016). For example, two cyanobacterial viruses (LBVG_2 and LBVG_12) harbored the photosystem II D1 protein (*psbA*) gene, which accelerates the host’s photosynthetic light reaction rates to facilitate viral propagation. LBVG_55, one of the dominant viruses in the hypolimnion with an unknown host (Fig. 5), encoded genes (*cobS* and *cobT*) involved in the biosynthesis of cobalamin, which is often absent from bacterial genomes, thus limiting their metabolic capability (Shelton *et al.*, 2019). Several LBVGs harbored a cluster of genes involved in the biosynthesis of lipopolysaccharides (LPS) and capsular polysaccharides. For example, LBVG_26 encoded genes involved in the biosynthesis of N-acetylneuraminic acid (*neuA*, *neuB*, and *neuC*), in addition to genes annotated as encoding putative sugar epimerase (*capD*), amidotransferase (*pseA*), aminotransferase (*pglC*), and acyltransferase (*pglD*). These genes may alter the cell surface properties of their hosts to prevent infection with other viruses (Sullivan *et al.*, 2005) or to provide tolerance to temperature, oxidative stress, or antibiotics (Nishimura *et al.*, 2017a). Proteins in the adenylyl-sulfate kinase (CysC) and 3’-phosphoadenosine 5’-phosphosulfate (PAPS) reductase (CysH) families were encoded in 4 and 17 LBVGs, respectively, and LBVG_98 contained both (Supplementary Dataset S1). These genes are involved in assimilatory sulfate reduction and likely facilitate the host’s utilization of reduced sulfur compounds (Summer *et al.*, 2007; Mizuno *et al.*, 2013a). These genes are present in ubiquitous and abundant viral groups (e.g., gOTU_3 and gOTU_11), indicating that they are quantitatively significant in the ecosystem. By contrast, these genes were not reported as common AMGs in marine ecosystems (Hurwitz *et al.*, 2015; Hurwitz and U’Ren, 2016). Given that freshwater systems are sulfur-limited, and that the genomes of many predominant freshwater bacterioplankton lineages such as LD12 (Eiler *et al.*, 2016; Henson *et al.*, 2018), acI-B1 (Neuenschwander *et al.*, 2018), and CL500-11 (Mehrshad *et al.*, 2018) lack the complete pathway for assimilatory sulfate reduction, these viral-encoded genes are likely key factors modulating microbial sulfur metabolism in the lake.

### Ecological role of viruses infecting the dominant bacterioplankton

Leveraged by the simultaneously reconstructed bacterioplankton genomes (LBMAGs), hosts were predicted for 40 LBVGs. The predicted hosts spanned 10 phyla (or Proteobacteria classes; Fig. 3), and the host of 6 LBVGs could be further resolved to the genus level (Table S5). Notably, we identified 13 LBVGs infecting Actinobacteria, which is one of the most diverse, abundant, and ubiquitous groups of freshwater bacterioplankton (Newton *et al.*, 2011). Aside from LBVG_68 (genome size = 40.0 kb), all actinoviral LBVGs had small genomes (14.0–19.8 kb) and formed a monophyletic clade composed of four gOTUs (gOTU_8, gOTU_13, gOTU_14, and gOTU_28; Fig. 2). Their monophyly was further supported by the proteomic tree including EVGs (https://www.genome.jp/viptree/u/LBV/retree/LBVGandEVG). All of them were closely related to viral genomes of the same size that originated from other freshwater systems (Fig. 6 and Supplementary Alignment S1). The members of gOTU_28 were relatives of the G4 actinophage (uvFW-CGR-AMDFOS-S50-C341) (Fig. 6), one of four actinoviral groups revealed metagenomically in a Spanish reservoir (Ghai *et al.*, 2017). Relatives of the other three Spanish actinoviral groups (G1, G2, and G3) were absent from our LBVGs but were observed among LBVCs (Fig. S7) and EVGs recovered from other freshwater lakes (Supplementary Alignment S2–4). Together, the results suggest that these actinoviral groups are ubiquitous in freshwater systems. Their broad range of GC contents (41.6–62.6%) suggests that their hosts include diverse members of Actinobacteria that also exhibit a wide range of GC contents (Table S3). In the epilimnion, LBVG_179 consecutively predominated and other actinoviral members also abundant in both the cellular and virion fractions (Figs. 5 and S8). Although data after September were not available in the epilimnion, the actinoviral predominance in the virion fraction after the onset of the mixing period (i.e., in the hypolimnion on January) implies that they were continuously dominated in the epilimnetic water until the end of stratification period. These results suggest that diverse members of actinovirus actively replicating and lysing the host cells to produce virions in the epilimnion. This finding is intriguing given the predominance of Actinobacteria and evokes the “King of the Mountain” (KoM) hypothesis, which was proposed to explain the co-dominance of marine *Pelagibacter* and its viruses (Giovannoni *et al.*, 2013). In the KoM hypothesis, the predominance of the host species under high viral lytic pressure provides a greater chance for the host to acquire resistance against viral infection through genomic recombination, and this generates a positive feedback loop for host propagation. Supporting this hypothesis, single-cell amplified genomes of acI, the most representative freshwater Actinobacterial lineage, exhibited high intra-population diversification, suggesting that acI members have a high recombination rate (Garcia *et al.*, 2018). Another study revealed that members of the acI lineage commonly harbor genomic islands containing genes that may be involved in viral host recognition (e.g., extracellular structural genes), even among closely related strains (average nucleotide identity [ANI] >97%) inhabiting the same lake (Neuenschwander *et al.*, 2018). Accordingly, the most closely related pair of actinoviral LBVGs in the present study (LVBG_169 and LBVG_172; *S*_G_ = 0.91, ANI = 96.4%) showed diversification in tail-related genes (e.g., tape measure protein), which are often involved in viral recognition of the host cell surface (Mizuno *et al.*, 2014; Rodriguez-Valera *et al.*, 2014) (Supplementary Alignment S1). Moreover, the streamlined acI genomes do not harbor the CRISPR-Cas system (Neuenschwander *et al.*, 2018), and a recent meta-epigenomic analysis conducted at Lake Biwa demonstrated that the genomes of Actinobacteria lack both methylated DNA motifs and methyltransferase genes (Hiraoka *et al.*, 2019), implying that they lack the restriction-modification system for resisting viral infection. These findings further support the KoM hypothesis, which assumes that the host prefers a strategy to overcome resource competition rather than act as a defense specialist (Giovannoni *et al.*, 2013). In light of this hypothesis, the high abundance and diversity of actinoviruses may result from an arms race with their hosts, leading to genomic diversification of the hosts to avoid mortality induced by a high viral load. It should be noted that such microdiversification of viral genomes poses challenges for metagenomic assembly (Martinez-Hernandez *et al.*, 2017; Roux *et al.*, 2017b; Warwick-Dugdale *et al.*, 2019); indeed, 109 LBVCs encoded the actinoviral hallmark *whiB* gene (Supplementary Dataset S1), suggesting that the majority of actinoviral genomes were not recovered as LBVGs (i.e., circularly assembled). Although further investigation is beyond the scope of the present study, our results revealed that, analogous to their hosts, actinoviruses are one of the most diverse, abundant, ubiquitous, and active viral groups in freshwater systems.

**Figure 6.**
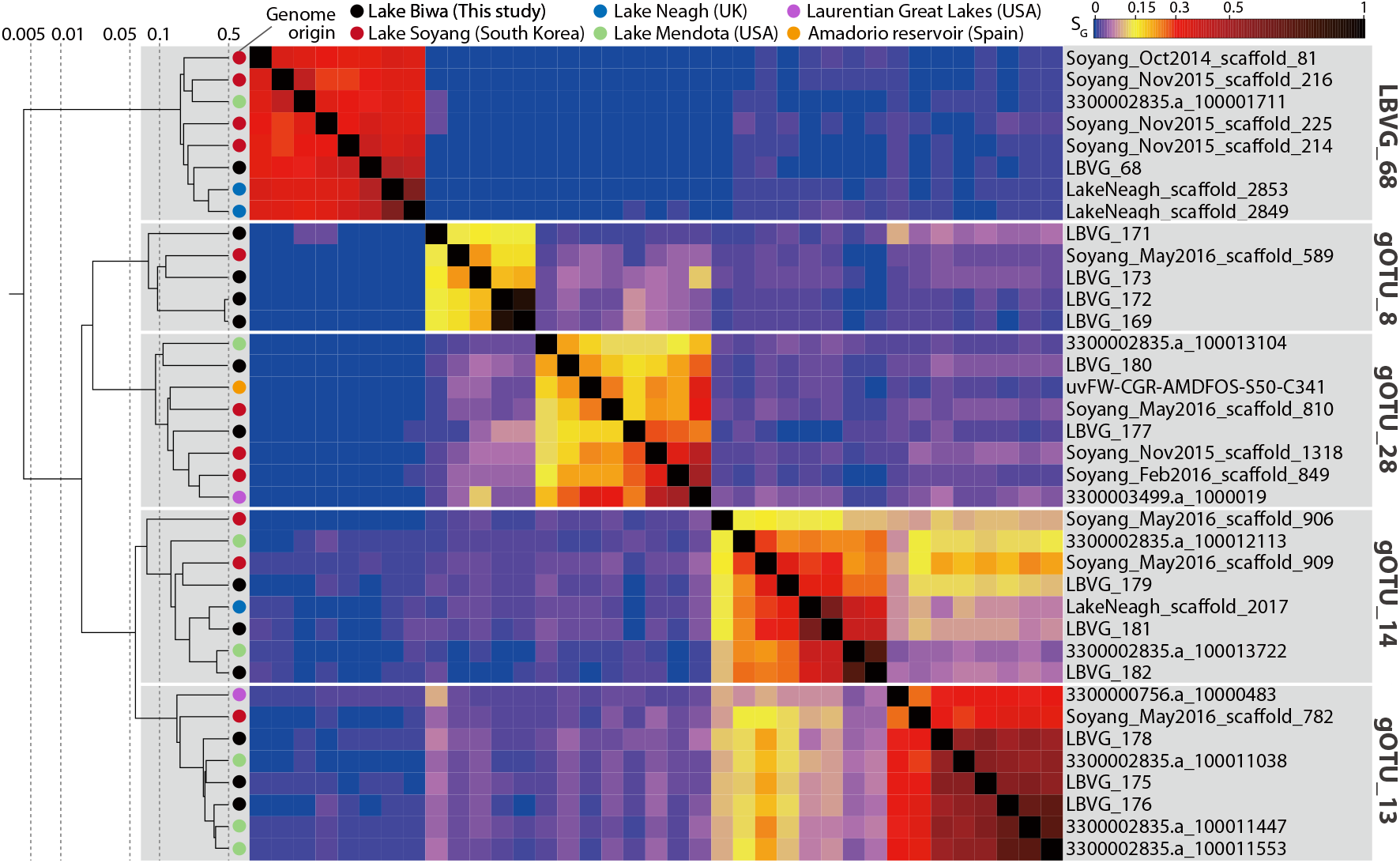
Proteomic tree and *S*_G_-based pairwise distance matrix among five actinoviral groups identified in the present study and their close relatives in the Environmental Viral Genome (EVG) database. Alignments between individual genomes can be found in Supplementary Alignment S1.

LD12 (*Ca*. Fonsibacter) is a freshwater division of the SAR11 lineage (Pelagibacteraceae), which is one of the most ubiquitous and abundant bacterioplankton in freshwater, as is the case for its marine counterparts (Salcher *et al.*, 2011; Henson *et al.*, 2018). Indeed, LD12 was by far the most abundant LBMAG in the epilimnion throughout the study period (Table S4). Unlike actinoviruses, the three predicted LD12 viruses were not clustered with other LBVGs (Fig. 2) and thus their genomic diversification seems limited, possibly reflecting the fact that LD12 bacteria also show limited genomic diversification (Zaremba-Niedzwiedzka *et al.*, 2013; Garcia *et al.*, 2018). Among these LBVGs, LBVG_1 was the largest (317.9 kb) viral genome assembled in the present study. LBVG_1 encoded many genes closely related to those of Pelagibacterales, including the 30S ribosomal protein S21, which is the most abundant virus-encoded ribosomal protein in aquatic systems and presumably facilitates takeover of the host’s translational machinery (Mizuno *et al.*, 2019) (Supplementary Dataset S1). A BLASTp search against the NCBI RefSeq database revealed that LBVG_1 showed high similarity to another large (genome size = 490 kb) alphaproteobacterial (Agrobacterium) phage Atu_ph07 (Attai *et al.*, 2018) in their jumbo phage marker genes (Yuan and Gao, 2017): terminase large subunit (amino acid sequence identity = 52.6%), major capsid protein (58.3%), and DNA polymerase family B (43.8%). These results suggest that LBVG_1 is a jumbo phage infecting the LD12 lineage. LBVG_1 was abundant in the cellular fraction in the epilimnion but rare in the virion fraction (Figs. 5 and S8). However, it is possible that their planktonic virions could have been included in the cellular fraction, because large virions of jumbo phage may be captured by the 0.22 µm pore-sized filter (Yuan and Gao, 2017). By contrast, the two other predicted LD12 viruses (LBVG_76 and LBVG_174) were abundant in both the cellular and virion fractions (Figs. 5 and S8), suggesting that these viruses were actively replicating and lysing their hosts. These results indicate that, in accordance with their marine counterparts, viruses are an important driving factor for the population dynamics of LD12, which is one of the most quantitatively significant components of freshwater microbial ecosystems.

The present study recovered a LBMAG of *Ca*. Nitrosoarchaeum (Fig. 3), which is a member of the ammonia-oxidizing, chemoautotrophic archaeal taxon Marine Group I (MGI). Members of MGI dominate the hypolimnion of deep freshwater lakes (Okazaki *et al.*, 2017) (Table S4) and thus are important players in the nitrogen and carbon cycles of lakes. In marine systems, members of MGI are also dominant in the deep aphotic layer, and recent studies have reported the presence of several putative MGI viruses (Chow *et al.*, 2015; Labonté *et al.*, 2015), including those encoding AMGs such as *amoC* (Ahlgren *et al.*, 2019) and *cobS* (López-Pérez *et al.*, 2018). The present study predicted one *Ca*. Nitrosoarchaeum virus (LBVG_51), the first freshwater MGI virus to our knowledge. The virus did not show genomic similarities to known marine MGI viruses or to any genomes in the EVG database. The genome encoded RadA-like ATPase, replication protein A and minichromosome-maintenance helicase genes, which drive the archaeal DNA replication system (Zatopek *et al.*, 2018), presumably to facilitate viral genomic replication in the host (Krupovic *et al.*, 2018). LBVG_51 was mainly detected in the cellular fraction in the hypolimnion (Figs. 5 and S8), suggesting that they were intracellular during the study period. While the relative abundances of the virus (LBVG_51) and the host (Ca. Nitrosoarchaeum) both showed a continuously increasing trend in the hypolimnion during the stratification period, the viral abundance increased disproportionally faster (48.3-fold from September to December) than that of the host (2.4-fold) (Supplementary Dataset S1 and Table S4). No integrase gene was annotated in the LBVG_51 genome. Collectively, we hypothesize that LBVG_51 genomes were replicating extrachromosomally in cells of the host using the replication machinery encoded on the viral genome. A previous study reported an abrupt drop in *Ca*. Nitrosoarchaeum abundance after the onset of the mixing period (Okazaki and Nakano, 2016), but no evidence for mass induction was found in the present study. We further speculated that one incomplete contig, LVBC_1935 (16,737 bp), might also be derived from a *Ca*. Nitrosoarchaeum virus, as it encoded multiple archaeal genes and exhibited a GC content and spatiotemporal distribution corresponding to those of *Ca*. Nitrosoarchaeum (Supplementary Dataset S1).

Viruses infecting other bacterioplankton taxa, namely Cyanobacteria, Bacteroidetes, Betaproteobacteria, Gammaproteobacteria, Nitrospira, Planctomycetes, and Verrucomicrobia were also predicted (Figs. 2 and 3). Most of these virus-host pairs were novel, and their potential interaction with hosts is discussed in Supplementary Results and Discussion.

### Viral genomic diversification among habitats

Majority (116 of 183) of the LBVGs were closely related to genomes that were recovered from other freshwater habitats (Fig. 2), indicating that they belonged to ubiquitous freshwater viral lineages. Some of these lineages exhibited remarkably high inter-lake genomic similarities. For example, among actinoviruses (Fig. 6), LBVG_182 and 3300002835.a_100013722, a genome retrieved from Lake ^Mendota, USA (Roux *et al.*, 2017a), exhibited an *S*_G_ value of 0.76 and an ANI value of 84%^ (Supplementary Alignment S1). Moreover, LBVG_144 and Soyang_Oct2014_scaffold_160, which ^has an unknown host, exhibited an *S*_G_ value of 0.94 and an ANI value of 99% (Supplementary^ Alignment S5). Whereas nearly identical viral genomes have been reported from distant locations in marine environments, such as the Mediterranean Sea and Pacific Ocean (Rodriguez-Valera *et al.*, 2014), these results in freshwater systems are intriguing, as lakes and reservoirs are physically isolated from one another. By contrast, the genomes of their host (i.e., bacterioplankton) inhabiting different freshwater systems are generally distinct (Hahn *et al.*, 2016; Kang *et al.*, 2017; Garcia *et al.*, 2018; Neuenschwander *et al.*, 2018), although highly similar genomes (ANI >95%) have occasionally been reported from lakes on different continents (Mehrshad *et al.*, 2018). The high phylogeographic connectivity of viruses was presumably because viruses can easily colonize a new habitat due to the lack of immunity by local hosts, whereas colonization by bacteria is often constrained by the priority effect, that is, occupation of the niche by prior inhabitants limits settlement of newly migrated genotypes (Incagnone *et al.*, 2015).

Aside from this evidence of connectivity among freshwater habitats, we also observed 34 ^LBVGs that were similar (*S*_G_ >0.15) to marine viral genomes (Fig. 2). For example, gOTU_26^ (LBVG_135 and LBVG_162)— one of the dominant viruses in the hypolimnion with an unidentified host (Fig. 5)— had a conserved genomic structure with viral genomes originating from a broad range of aquatic habitats; surface and deep waters of the Mediterranean Sea, Pacific Ocean, Atlantic Ocean, Arabian Sea, Osaka Bay, Lake Neagh and Lake Soyang (Supplementary Alignment S6). As observed for cyanoviruses (Dreher *et al.*, 2011; Chénard *et al.*, 2015), close phylogenetic relationships among marine and freshwater viruses suggest that their hosts are bacterioplankton belonging to taxa containing both marine and freshwater members, such as Pelagibacteriaceae (Eiler *et al.*, 2016), Methylophilaceae (Salcher *et al.*, 2015), and Flavobacteriaceae (Eiler and Bertilsson, 2007). Indeed, LBVG_76 and LBVG_174, which are putative LD12 viruses, share genomic structures with marine SAR11 phages recovered from Mediterranean viromes (Mizuno *et al.*, 2013b, 2016) and the *Tara* Oceans Virome (Brum *et al.*, 2015) (Supplementary Alignments S7 and S8). A recent study (Paver *et al.*, 2018) has challenged the idea that marine-freshwater transitions by bacterioplankton are infrequent (Logares *et al.*, 2009; Eiler *et al.*, 2016), and viral diversification processes that allow crossing of this salt divide in association with their hosts are an intriguing research topic. Notably, the greatest similarity between marine and freshwater viral genomes was observed among members of the *Synechococcus* viruses S-EIV1, LBVG_9 and TARA_ERS488929_N000037 (collected at a pelagic site in the Arabian Sea), which exhibited higher similarity scores (*S*_G_ = 0.52) than among those collected in freshwater systems or even within the same lake (Supplementary Alignment S9). Thus, in the diversification processes of at least some viral lineages, the transition between marine and freshwater habitats appears to be a relatively frequent event, presumably because viruses require less modification in their genetic machinery to cross the salt divide.

## Experimental Procedures

### Sample collection, sequencing, and assembly

Details of the sample collection, sequencing, and assembly procedures are available in Supplementary Materials and Methods and Figure S1. Briefly, the cellular fraction (0.22–5.0 μm) was obtained shipboard through filtration of the water sample. The virion fraction (<0.22 μm) was collected from filtered water by concentrating virions using iron(III) chloride flocculation (John *et al.*, 2011), followed by purification using CsCl density gradient ultracentrifugation (Hurwitz *et al.*, 2013). After DNA extraction and library preparation for shotgun metagenomics, two sequencing runs (2 × 300 bp) were performed using the Illumina MiSeq system for each of the cellular and virion fractions. The sequence reads generated were (co-)assembled using metaSPAdes v. 3.10.1 (Nurk *et al.*, 2017) for the cellular fraction and SPAdes v. 3.9.0 (Bankevich *et al.*, 2012) for the virion fraction (Table S2).

### Extraction of viral contigs

Viral contigs were extracted from 11 (co-)assemblies from the virion fraction and 7 assemblies from the cellular fraction (Table S2). Only contigs longer than 10 kbp were considered for subsequent analysis. Identification of viral contigs was performed with VirSorter v. 1.0.3 (Roux *et al.*, 2015) using Virome Decontamination mode with reference to the Virome Database, and all contigs assigned to categories 1–3 (virus and prophage) were extracted. The resulting 8,173 contigs were inspected for redundancy using BLASTn searches against themselves with NCBI-BLAST+ v. 2.7.1 (Camacho *et al.*, 2009), where all >500 nt high-scoring segment pairs were considered, contigs with >95% nucleotide identity across >80% of the contig length (Roux *et al.*, 2017b) were identified as redundant, and the shorter contig in the pair was discarded. The resulting 4,315 contigs were further decontaminated through inspection for viral hallmark genes. To this end, 134,388 open reading frames (ORFs) on 4,315 contigs were predicted with Prodigal v. 2.6.3 (Hyatt *et al.*, 2010) and functionally annotated (see Supplementary Materials and Methods for functional annotation workflow). Genes annotated as encoding terminase, tail, capsid or portal proteins were regarded as viral hallmark genes. Among contigs with no viral hallmark genes, those annotated as prophages using VirSorter (i.e., containing cellular hallmark genes) and those exclusively detected in the cellular fraction based on read coverage were regarded as bacterial genomic contaminants and were removed. Consequently, 4,158 contigs (LBVCs) were retained. Further, 183 complete (i.e., circular) genomes (LBVGs) were identified using ccfind v. 1.1 (https://github.com/yosuken/ccfind) (Nishimura *et al.*, 2017a). Handling of FASTA files and calculations of genome size and GC contents were performed using SeqKit v. 0.5.0 (Shen *et al.*, 2016).

### Abundance estimation based on metagenomic read coverage

The relative abundance (as metagenomic read coverage shown by FPKM) of LBVGs in each sample was calculated using the CountMappedReads v. 1.0 script (https://github.com/yosuken/CountMappedReads). Habitat preference for the epilimnion over the hypolimnion (*P*_epi_) was measured as the quotient of abundance in the epilimnion versus the sum of abundance in the epilimnion and hypolimnion (i.e., epilimnion/[epilimnion + hypolimnion]), and habitat preference for the cellular fraction over the virion fraction (*P*_cell_) was measured as the quotient of abundance in the cellular fraction versus the sum of abundance in the cellular and virion fractions (i.e., cellular/[cellular + virion]). Note that only samples for which both virion and cellular fractions were available (i.e., July–September in the epilimnion and September–December in the hypolimnion) were considered for this calculation.

### Binning of prokaryotic genomes

Binning of prokaryotic genomes was performed with MetaBAT v. 0.32.5 (Kang *et al.*, 2015) using the (co-) assemblies from the cellular fraction, and the output was manually curated (Fig. S1 and Table S2; see Supplementary Materials and Methods for details). Phylogenetic assignment of the resulting 57 bins (LBMAGs) was carried out by drawing a phylogenetic tree with published freshwater bacterioplankton genomes using PhyloPhlAn v. 0.99 (Segata *et al.*, 2013) and the assignments were confirmed using GTDB-Tk v. 0.2.1 with reference to the release 86 database (Parks *et al.*, 2018). The relative abundance of LBMAGs in each sample was calculated using the CountMappedReads v. 1.0 script.

### Construction of reference databases

The RVG and EVG databases were originally described by Nishimura et al. (2017a) and were updated in the present study (Fig. S2). The RVG was prepared by extracting 2,621 genomes of isolated dsDNA prokaryotic viruses from the Virus-Host Database (as of Feb. 6, 2018) (Mihara *et al.*, 2016). The EVG was built based on 1,811 genomes compiled for the previous study (Nishimura *et al.*, 2017a) and augmented with the following datasets: (i) 125,842 mVCs (metagenomic Viral Contigs), which were generated from metagenomes deposited in the IMG/M database (Paez-Espino *et al.*, 2016); (ii) 28 complete viral genomes retrieved from a Mediterranean deep-ocean virome (Mizuno *et al.*, 2016); (iii) contigs assembled from six viromes collected seasonally from Lake Soyang (PRJEB15535) (Moon *et al.*, 2017); (iv) 488 contigs previously compiled from published freshwater viromes (Arkhipova *et al.*, 2018); and (v) 313 complete viral genomes recovered from Lake Neagh (Arkhipova *et al.*, 2018). In addition, the genomes of recently reported freshwater viruses, including the *Synechococcus* virus S-EIV1 (Chénard *et al.*, 2015), eight actinoviruses (Ghai *et al.*, 2017), a LD28 virus (Moon *et al.*, 2017), two Comamonadaceae viruses (Moon *et al.*, 2018), and a *Polynucleobacter* virus (Cabello-Yeves *et al.*, 2017) were compiled. Assembly of the Lake Soyang virome was performed using the same parameters used for the Lake Biwa virome. The contigs were inspected for circularity using ccfind v. 1.1. Only >10 kb complete genomes were retained and dereplicated using the BLASTn search against themselves, where all >500 nt high-scoring segment pairs were considered and those with >99% nucleotide identity across >95% of their length were identified as duplicates. The resulting EVG database included 2,860 viral genomes.

### Phylogenetic analysis of viral genomes

Calculation of *S*_G_ and construction of a *S*_G_-based proteomic tree were performed using ViPTreeGen v. 1.1.1, a standalone version of ViPTree (Nishimura *et al.*, 2017b). LBVGs were searched against the RVG and EVG databases using the 2D mode of ViPTreeGen v. 1.1.1, and the number of hits with *S*_G_ >0.15 was determined for each LBVG.

### Host prediction

The host of each LBVG was predicted from combinations of *in silico* analyses (Edwards *et al.*, 2016). Briefly, the host of an LBVG was predicted when a close relative infected a known host, a taxon-specific marker gene was found, >80% of annotated bacterial genes were taxonomically related to a single taxon, or the virus and host genomes shared a >30 bp nucleotide sequence segment. See Supplementary Materials and Methods for a detailed workflow.

### Data availability

Raw reads from the cellular and virion fractions were deposited under accession numbers PRJDB6644 and PRJDB7309, respectively. LBMAGs were deposited under the BioSample identifiers SAMD00166046–00166102. The nucleotide sequences of the LBVGs, LBVCs, RVG, and EVG are available from https://doi.org/10.6084/m9.figshare.7934924.

## Acknowledgments

This work was supported by JSPS KAKENHI (Nos. 15J00971, 16H06429, 16K21723, 16H06437, 17H03850, 17K19289, and 18J00300), the Environment Research and Technology Development Fund (No. 5-1607) of the Ministry of the Environment, Japan, and the Canon Foundation (No. 203143100025). Computation time was provided by the SuperComputer System, Institute for Chemical Research, Kyoto University. Field sampling on Lake Biwa was conducted using the research vessel Hasu belonging to the Center for Ecological Research, Kyoto University, and supported by Y. Goda and T. Akatsuka. We are grateful to T. Narihiro and M. K. Nobu for their helpful advice about metagenomic analysis, and to H. Watai, N. Haruki, and M. Yamanaka for their assistance in sample preparation. We thank K. Arkhipova and K. Moon for providing the original datasets from their published works.

