## Supplementary_Information for "Genome-resolved viral and cellular metagenomes revealed potential key virus-host interactions in a deep freshwater lake"

**This PDF file includes:**

Supplementary Materials and Methods
Supplementary Results and Discussion
Figures. S1 to S9
Captions for Tables S1 to S6
Captions for Dataset S1

**Other supplementary materials for this manuscript include the following:**

Tables S1 to S6
Dataset S1

### Supplementary Materials and Methods

#### *Sample collection, sequencing, and assembly*

Water samples were collected monthly from the epilimnion (5 m) and hypolimnion (65 m) at a pelagic station (35°13′09.5″N, 135°59′44.7″E) at Lake Biwa, Japan (water depth: ca. 73 m) from June 2016 to February 2017 (Table S1, Fig. S1). Vertical profiles of water temperature and dissolved oxygen were determined *in situ* using a CTD profiler (SBE-911plus; Sea Bird Electronics). Water sample filtration was performed using an onboard peristaltic pump within 1 h after sample collection: After prefiltration using a 5.0 µm pore polycarbonate filter (Whatman, cat. no. 111113), the cellular fraction was collected on a 0.22 µm pore polyethersulfone filter cartridge (Millipore, Sterivex SVGP01050) until the Sterivex filter clogged (1–2 and 2–3 L for the epilimnetic and hypolimnetic samples, respectively). The sampled Sterivex cartridges were stored at –20°C until DNA extraction. To collect the virion fraction, virions in 2 L filtered (<0.22 µm) water were concentrated using the $FeCl_3$ flocculation method (John *et al.*, 2011) and purified by CsCl density ultracentrifugation targeting densities of 1.45–1.5 g $mL^{-1}$ (Hurwitz *et al.*, 2013). Extracted virions were stored in modified SM buffer (50 mM Tris-HCl, pH 7.5; 100 mM NaCl; 10 mM $MgSO_4$) by replacing the CsCl solution using a dialysis membrane and kept in the dark at 4°C until DNA extraction. We fixed 5 mL water samples with 1% glutaraldehyde for epifluorescence microscopic enumeration of prokaryotic cells (Porter and Feig, 1980) and virus-like particles (Noble and Fuhrman, 1998). All chemical solutions were prefiltered using a 0.02 µm syringe filter (Anotop, Whatman) before use to remove potential contaminants.

To extract DNA from the cellular fraction, the filter paper was removed from the Sterivex cartridge, chopped into small pieces (ca. 3 × 3 mm) using a sterile scalpel, and processed using the PowerSoil DNA Isolation Kit (MoBio Laboratories) following the manufacturer’s instructions. At least 585 ng DNA was obtained from each sample. Extracted DNA was sheared to 800 bp on average using an ultrasonicator (Covaris), and polymerase chain reaction (PCR)-free sequencing libraries were prepared using the KAPA Hyper Prep Kit (Kapa Biosystems) following the manufacturer’s instructions. Three samples (July–September) from the epilimnion and four samples (September–December) from the hypolimnion were equimolarly pooled and sequenced (i.e., two runs in total) using the Illumina MiSeq platform with v3 chemistry (2 × 300 bp).

DNA extraction from the virion resuspension was performed using the xanthogenate method (Tillett and Neilan, 2000) with the initial DNase (Invitrogen, TURBO DNase) decontamination step as described in a previous study (Nishimura *et al.*, 2017). Sequencing libraries were prepared using the Nextera XT DNA sample preparation kit (Illumina) according to the manufacturer’s instructions, except that 0.25 ng DNA was used as input (Nishimura *et al.*, 2017). Two sequencing runs were performed using the Illumina MiSeq platform with v3 chemistry (2 × 300 bp) as follows: six samples occupied 1/4 of one run generating > 9 million reads, and the other six samples occupied 1/12 of one run generating > 3 million reads (Table S1).

Reads from the cellular fraction were inspected by FastQC (http://www.bioinformatics.babraham.ac.uk/projects/fastqc/) and the identified artificial sequences (e.g., adaptor readthrough) were removed using Trimmomatic v. 0.36 (Bolger *et al.*, 2014), followed by quality trimming using PRINSEQ lite v. 0.20.4 (Schmieder and Edwards, 2011) with the following parameters: trim_qual_right: 20, min_len: 50, and trim_qual_window: 6. The resulting high-quality reads from each sample were assembled using metaSPAdes v. 3.10.1 (Nurk *et al.*, 2017) with k-mer lengths of 21, 33, 55, 77, 99, and 127. Co-assemblies of pooled reads from the three epilimnetic samples and four hypolimnetic samples were performed using the same parameters (Table S2).

Reads from the virion fraction were directly assembled by SPAdes v. 3.9.0 (Bankevich *et al.*, 2012) using the '--careful' option and k-mer lengths of 21, 33, 55, 77, 99, and 127, following the optimized genome assembly strategy proposed by a previous study (Nishimura *et al.*, 2017). The six deeply sequenced samples were assembled individually. Co-assembles were performed using combinations of the samples including those that were shallowly sequenced (Table S2).

#### *Workflow for gene functional annotation*

Open reading frames (ORFs) in the 4,315 contigs after the dereplication step (Fig. S1) were predicted by Prodigal v. 2.6.3 (Hyatt *et al.*, 2010) using the '-p meta' option. The resulting 134,388 amino acid sequences were annotated by eggNOG-Mapper v. 1.0.3 (Huerta-Cepas *et al.*, 2017) using the '-m diamond' option with a seed ortholog e-value cutoff of $10^{-5}$ for general annotation, and by the hmmscan search in the HMMER v. 3.1b2 software (Eddy, 2011) against Prokaryotic Virus Orthologous Groups (pVOGs) hidden Marcov model (HMM) profiles (Grazziotin *et al.*, 2017) with an e-value cutoff of $10^{-5}$ for virus-specific annotation. For each of the amino acid sequences, a similarity search against the UniRef90 (release 2018_01) database (Suzek *et al.*, 2007) was performed using DIAMOND v. 0.9.14 (Buchfink *et al.*, 2015) with an e-value cutoff of $10^{-5}$. A sensitive HMM–HMM search against the Pfam v. 31.0 database (Finn *et al.*, 2016) was also performed using HHsearch (Söding, 2005) and JackHMMER (Johnson *et al.*, 2010). For this purpose, a reference of viral proteins containing 3,144,341 amino acid sequences derived from the reference viral genome (RVG), the original EVG by Nishimura *et al.* (2017), and the original mVCs (Paez-Espino *et al.*, 2016) was created by predicting ORFs using Prodigal v. 2.6.3 (Hyatt *et al.*, 2010) with the '-p meta' option. Subsequently, the 134,388 amino acid sequences were iteratively searched against the 3,144,341 reference sequences using JackHMMER to generate query HMMs. Each query HMM was searched against the Pfam database and the hit with the highest probability score with a cutoff probability score of 95% was determined for each query. This procedure was performed using pipeline_for_high_sensitive_domain_search v. 0.1.0 (https://github.com/yosuken/pipeline_for_high_sensitive_domain_search) (Nishimura *et al.*, 2017; Yoshida *et al.*, 2018). Functional annotation results for each pipeline are available in Supplementary Dataset S1.

#### *Workflow for prokaryotic genome binning*

The seven individual assemblies and two co-assemblies from the epilimnion and hypolimnion (Table S2) were used for metagenomic binning by MetaBAT v. 0.32.5 (Kang *et al.*, 2015). For coverage calculation, the high-quality reads (those used for assembly) from the cellular fractions were mapped against the assembled contigs using the BWA-MEM algorithm of BWA v. 0.7.15 (Li and Durbin, 2009), and the resulting sam files were transformed to bam files using Samtools v. 0.3.1 (Li *et al.*, 2009). Binning was performed separately for the epilimnetic and hypolimnetic samples, i.e., cross-mapping for coverage calculation was performed among months but not between water layers. MetaBAT was run using the '-m 1500' and '-B 20' options. Quality of the resulting bins was inspected using checkM v. 1.0.7 (Parks *et al.*, 2015) with default parameters, and bins with >50% completeness and <30% contamination were retained. Because the bins were derived from multiple samples in the same lake, they contained redundancy. For dereplication, all generated bins were clustered by generating a genome-resolved phylogenetic tree using

PhyloPhlAn v. 0.99 (Segata *et al.*, 2013). Integrity of the clusters was inspected using MUMmer v. 3.23 (Kurtz *et al.*, 2004) and bins within the same cluster were confirmed to share >95% (>99% in most cases) nucleotide identity. Subsequently, a representative bin was selected for each cluster as that with the highest completeness and <5% contamination; if such a bin did not exist, then the least contaminated bin was selected. Singleton bins that did not form a cluster in the tree (i.e., failure to replicate in another assembly) were discarded to ensure bin robustness. The resulting 57 bins were designated as LBMAGs (Lake Biwa Metagenome Assembled Genomes).

#### *Workflow for host prediction.*

The host of an LBVG (Lake Biwa Viral Genome) was predicted when a close relative infected a known host, a taxon-specific marker gene was found, >80% of annotated bacterial genes were taxonomically related to a single taxon, or the virus and host genomes shared a >30 bp nucleotide sequence segment. The workflow for the prediction is summarized below and in Table S5.

***A close relative infected a known host.***

If a given LBVG was closely related ($S_G > 0.15$) to a viral genome whose host is known, the LBVG was predicted to share the host with the reference at the phylum level. By this criterion, the hosts of four LBVGs (LBVG_2, _65, _81, and _180) were predicted as they were closely related to described viral genomes (i.e., RVG). The hosts of an additional nine LBVGs were predicted as they were closely related to EVGs whose host was previously identified or predicted (Table S5). Among these, the hosts of LBVG_76 and LBVG_84 were further identified at a finer taxonomic resolution, as LD12 (*Ca.* Fonsibacter) and LD28 (*Ca.* Methylopumilus), because their close relatives were a putative pelagiphage (Mizuno *et al.*, 2013) and an isolated LD28 virus (Moon *et al.*, 2017), respectively.

***A taxon-specific marker gene was found.***

LBVG_68 and LBVG_180 harbored the actinobacterial transcriptional factor *whiB*, suggesting that their hosts are Actinobacteria (Ghai *et al.*, 2017). LBVG_51 had the RadA-like ATPase, replication protein A (RPA), and minichromosome maintenance (MCM) helicase genes, which drive the archaeal DNA replication system (Zatopek *et al.*, 2018) and are often encoded in known archaeal viruses (Krupovic *et al.*, 2018), indicating that the host is archaeal. In Lake Biwa, *Ca.* Nitrosoarchaeum of the MGI group was the only archaeal lineage in the water column reported by previous studies (Okazaki and Nakano, 2016; Okazaki *et al.*, 2017) and indeed it was the sole archaeal LBMAG reconstructed in the present study (Fig. 3); therefore, the host of LBVG_51 was predicted as *Ca.* Nitrosoarchaeum.

***>80% of the annotated bacterial genes were taxonomically related to a single taxon.***

For each gene of the LBVGs, the phylogenetic affiliation of the closest protein sequence in the UniRef90 database (DIAMOND search; e-value cutoff: $10^{-5}$) was determined. If a single phylum (or class for Proteobacteria) occupied >80% of the affiliations of the annotated bacterial genes in a viral genome, the phylum was predicted as host of the virus. Using this criterion, four Bacteroidetes (LBVG_79, _120, _126, and _183) and four Actinobacteria (LBVG_171, _172, _173, and _181) viruses were predicted.

***The virus and host genomes shared a >30 bp nucleotide sequence segment.***

If a shared nucleotide sequence segment (>30 bp, allowing up to one mismatch) was found between a viral and a bacterial genome, they were considered a virus–host pair. To this end, all LBVG sequences were searched by BLASTn against a set of freshwater bacterioplankton genomes, including the 57 LBMAGs and published bacterial genomes listed in Figure S3. Given the relatively high false positive rate of prediction based on a short (ca. 30–60 bp) nucleotide sequence match (Edwards *et al.*, 2016), predictions were collectively supported by further information, considering whether (i) there was an adjacent integrase gene, which indicates that the shared sequence is the phage attachment site (attP), (ii) the shared sequence is a part of a tRNA sequence, where the attP is often located, and (iii) the shared sequence is encoded in a CRISPR spacer, which indicates that the host acquired immunity against the virus in response to a recent infection event (Edwards *et al.*, 2016). Host CRISPR spacers and tRNA sequences were determined by metaCRT (Bland *et al.*, 2007; Rho *et al.*, 2012) and ARAGORN v. 1.2.36 (Laslett and Canback, 2004), respectively. Using these criteria, hosts were predicted for six LBVGs (Table S5). Among these, sequences shared by four LBVGs (LBVG_24, _36, _96, and _174) were located adjacent to an integrase gene, and two of these (LBVG_36 and _96) were located within a tRNA sequence. LBVG_96 shared its partial tRNA sequence with LD28 and Limnohabitans of both LBMAG and the published genome; thus, their host was predicted at the class level, as Betaproteobacteria. The host of LBVG_102 was predicted by exact sequence matches to CRISPR spacer sequences in the *Methylobacter* LBMAG. The host of LBVG_60 was predicted because the virus shared not only two 40 bp exact matches but also six long nucleotide segments (2195–7177 bp, >98% identity) with the LBMAG Verrucomicrobia_hypo.bin2. The predictions for LBVG_36, _60, and _174 were further supported by close genomic 6-mer composition between the virus and host, calculated by VirHostMatcher v. 1.0.0 (Ahlgren *et al.*, 2017) using the default parameters (Table S6).

We also predicted the host of the following LBVGs by collectively considering multiple types of evidence: LBVG_1 was predicted as an LD12 (*Ca.* Fonsibacter) virus, because it shares an 866 bp nucleotide segment (77.0% identity) with the *Ca.* Fonsibacter ubiquis LSUCC0530 genome (CP024034), and 120 genes of LBVG_1 (>50% of the annotated genes) were closely related to Alphaproteobacteria genes, of which 49 were Pelagibacterales (including 30S ribosomal protein S21) and four were *Ca.* Fonsibacter (Supplementary Dataset S1). LBVG_175, _176, and _178 (gOTU_13) were predicted as actinovirus, because all of their viral hallmark genes (terminase, tail, capsid, and portal proteins) were closely related to those of known actinoviruses (Supplementary Dataset S1). The prediction was further supported by the viral proteomic tree (Fig. 2), which revealed that gOTU_13 is in a monophyletic cluster, in which all other members (i.e., gOTU_8, _14, and _28) were predicted as actinoviruses.

Finally, if the host is predicted for at least one members of a gOTU, all LVBGs in the gOTU (i.e., members sharing the $S_G > 0.15$ similarity) were predicted to share the host at the phylum level.

We note that the hosts of several other LBVGs can further be speculated, although they are not mentioned in the main text as they did not meet the prediction criteria described above. Members of gOTU_1 and gOTU_11 may be Betaproteobacteria viruses, because they had high numbers of genes (particularly viral structural genes) that were closely related to those of Betaproteobacteria or their viruses (Supplementary Dataset S1). LBVG_3, _4, _5, _6, and _7 may be Bacteroidetes viruses, because they had high numbers of genes that were closely related to the phylum (Supplementary Dataset S1).

### Supplementary Results and Discussion

#### *Virus–host interactions other than those mentioned in the main text*

***Cyanobacteria.***

Metagenomic binning recovered three cyanobacterial LBMAGs, all of which were affiliated with *Synechococcus/Cyanobium* sub-cluster 5.2 (Fig. S3), a cosmopolitan picocyanobacterial group found in freshwater, marine, brackish, and euryhaline systems (Cabello-Yeves *et al.*, 2018). Correspondingly, the present study recovered LBVGs closely related to cyanoviruses previously described from either marine or freshwater habitats, i.e., cyanomyovirus S-PM2 (a close relative of LBVG_2), cyanosiphovirus KBS-S-2A (LBVG_81), cyanovirus S-EIV1 (gOTU_10), and the metagenomically reconstructed putative cyanovirus uvMED-CGR-C15A-MedDCM-OCT-S31-C20 (Mizuno *et al.*, 2013) (gOTU_17) (Supplementary Dataset S1). Although these LBVGs were generally abundant in the epilimnion, following host habitat preference, they were occasionally also detected in the hypolimnion (Fig. S8), perhaps as a result of sinking cyanobacterial cells in the water column (Takasu *et al.*, 2015). In addition, 29 LBVCs (Lake Biwa Viral Contigs) encoded *psbA* or *psbD* (Supplementary Dataset S1), which are cyanoviral photosynthetic AMGs (Sullivan *et al.*, 2006), which implies the presence of a vast, unobserved diversity of cyanoviruses in the lake whose genome was not circularly assembled in the present study.

***Bacteroidetes.***

Six Bacteroidetes LBMAGs (Fig. 3) and five LBVGs predicted to infect Bacteroidetes (Fig. 2) were reconstructed in the present study. All five viruses had a close relative recovered from other freshwater habitats (Fig. 2), suggesting that their hosts are ubiquitous freshwater Bacteroidetes lineages (Newton *et al.*, 2011). Members of Bacteroidetes typically adopt an opportunistic or copiotrophic strategy, in which they grow quickly responding to a pulse of freshly supplied photosynthetic product from a phytoplankton bloom (Eiler and Bertilsson, 2007; Zeder *et al.*, 2009; Salcher, 2013). Accordingly, these viruses are likely responsible for rapid turnover of organic matter, in which fresh photosynthetic product incorporated into the host is instantly shunted back to the environment by lysis. Indeed, three Bacteroidetes viruses, LBVG_79, _126, and _148 each simultaneously exhibited a peak in abundance in both virion and cellular fractions of the epilimnion (Fig. S8), suggesting their rapid infection–replication–lysis cycle.

***Betaproteobacteria.***

Five Betaproteobacteria LBMAGs (Fig. 3) and seven predicted Betaproteobacteria viruses (Fig. 2) were reconstructed in the present study. One circular (LBVG_84) and one partial (LBVC_433) viral genome were closely related to P19250A (Moon *et al.*, 2017), a virus isolated from Lake Soyang (South Korea) infecting LD28 (*Ca.* Methylopumilus), a ubiquitous freshwater methylotrophic lineage (Salcher *et al.*, 2015) (Fig. S9). LBVG_84 was abundant in both cellular and virion fractions of the hypolimnion during the late stratification and mixing period (Figs. 5, S4 and S8), suggesting that the virus actively replicates and lyses the host in winter, as has also been proposed for A19250A in Lake Soyang (Moon *et al.*, 2017). Another study in Lake Soyang reported the isolation of two viral lineages infecting Comamonadaceae, a dominant freshwater Betaproteobacteria family (Moon *et al.*, 2018); however, their close relative was not detected in either LBVG or LBVC. *Polynucleobacter* is another predominant freshwater Betaproteobacteria genus (Hahn, 2003; Newton *et al.*, 2011). Previous studies have demonstrated its presence in Lake Biwa (Watanabe *et al.*, 2012; Okazaki and Nakano, 2016), and a putative *Polynucleobacter* virus has been reported from a Lake Baikal metagenome (Cabello-Yeves *et al.*, 2017b). However, no *Polynucleobacter* LBMAGs were recovered in the present study, and no evidence was found to support the presence of LBVG or LBVC of a *Polynucleobacter* virus.

***Gammaproteobacteria.***

An LBMAG of *Methylobacter*, a genus of typical methane oxidizing bacteria in freshwater systems (Borrel *et al.*, 2011), and a predicted virus infecting them (LBVG_102) were recovered in the present study. Both host (Table S4) and virus (Figs. S4 and S8) were dominant in the hypolimnion, consistent with a previous report that methane oxidizer is specific to aphotic hypolimnion (Murase and Sugimoto, 2005). Notably, this virus–host pair was the only case predicted by CRISPR spacer hits (Table S5). Accordingly, the abrupt drop in viral abundance observed in the cellular fraction in December (Fig. S8), which was coincident with relatively constant host abundance during the same period (Table S4), may have resulted from the host's viral immunity response.

***Nitrospira, Planctomycetes, and Verrucomicrobia*.**

The present study recovered an LBMAG of Nitrospira, a nitrifying bacteria dominant in the hypolimnia of deep freshwater lakes (Mukherjee *et al.*, 2016; Okazaki and Nakano, 2016; Okazaki *et al.*, 2017). Eight Planctomycetes and three Verrucomicrobia LBMAGs were also reconstructed (Fig. 3), supporting the recently proposed genome-resolved diversities of these phyla in freshwater systems (Cabello-Yeves *et al.*, 2017a; Andrei *et al.*, 2019). Most of these were preferentially distributed in the hypolimnion (Table S4). Correspondingly, the present study predicted viruses infecting the phyla Nitrospira (LVBG_36), Planctomycetes (LBVG_24), and Verrucomicrobia (LBVG_60) that were hypolimnion specialists (Fig. S8) without close relatives in the EVG database (Fig. 2). All of these encoded an integrase gene (Table S5), and LBVG_24 and LBVG_60 exhibited disproportionally higher abundance in the cellular fraction than in the virion fraction (Fig. S8), indicating their preference for the lysogenic state during the study period.

***The CL500–11 lineage (Chloroflexi).***

The hypolimnion-specific CL500–11 lineage (phylum Chloroflexi) is the most numerically and volumetrically significant bacterioplankton lineage in deep freshwater lakes (Okazaki *et al.*, 2013, 2017, 2018). Indeed, CL500–11 was by far the most abundant LBMAG in the hypolimnion (Table S4). However, no evidence confidently predicted an LBVG infecting CL500–11. We speculated that an incomplete contig, LBVC_2745 (13,561 bp), might be a genomic fragment of a CL500–11 virus, because it has viral structural genes closely related to those encoded in a published Chloroflexi genome and showed GC content and abundance dynamics corresponding to those of CL500–11 (Supplementary Dataset S1).

### *Supplementary proteomic tree and genomic alignment*

A proteomic tree including all Lake Biwa viral genomes (LBVGs) and environmental viral genomes (EVGs) (total: 3,043 genomes) is available on the ViPTree server (https://www.genome.jp/viptree/u/LBV/retree/LBVGandEVG). Alignments among the genomes and gene functional annotations can be accessed on the server by clicking a node on the tree. Detailed instructions are available in the ViPTree documentation (https://www.genome.jp/viptree/document). Among the tBLASTx alignments available on the server, the following (Supplementary Alignments S1–8) were selected and cited in the main text to support the discussion. The links below provide direct access to the alignment results. Note that each genome was reversed and/or circularly permutated for better alignment visibility. Unless otherwise stated, the genome order in each alignment follows that on the proteomic tree, such that the closest genomes were arranged to be adjacent.

**Supplementary Alignment S1**

Alignment of the five actinoviral groups identified in the present study and their close relatives found in the EVG database. Note that members and genome order correspond to those shown in Figure 6.

**Supplementary Alignment S2**

Alignment of the G1 actinovirus (KU886268) proposed by Ghai *et al.* (2017) and closely related genomes found in the EVG database.

**Supplementary Alignment S3**

Alignment of the G2 actinoviruses (KU886269, KU886271, and KU886273) proposed by Ghai *et al.* (2017) and related genomes found in the EVG database. Note that the three original G2 genomes were not closely related (i.e., $S_G > 0.15$) to each other, and the smallest monophyletic clade that included all three genomes also included a clade containing gOTU_68 (see original tree in https://www.genome.jp/viptree/u/LBV/retree/LBVGandEVG).

**Supplementary Alignment S4**

Alignment of the G3 actinoviruses (KU886270, KU886272, and KU886274) proposed by Ghai *et al.* (2017) and closely related genomes found in the EVG database.

**Supplementary Alignment S5**

Alignment of the viral genomic pair showing the highest inter-lake genomic similarity in the present study: LBVG_144 and Soyang_Oct2014_scaffold_160.

**Supplementary Alignment S6**

Alignment of gOTU_26 (LBVG_135 and 162) and closely related genomes found in the EVG database, including members originating from broad aquatic habitats: Surface (AP013516) and deep (KT997842) waters of the Mediterranean Sea, Pacific Ocean (TARA_ERS492198_N000332), Atlantic Ocean (TARA_ERS490285_N000586), Arabian Sea (TARA_ERS488813_N000325), Osaka Bay (OBV_N00119), Lake Neagh and Lake Soyang.

**Supplementary Alignment S7**

Alignment of LBVG_76 and closely related genomes found in the EVG database, which are putative pelagiphages recovered from surface (AP013442, AP013443) (Mizuno *et al.*, 2013) and deep (KT997865) (Mizuno *et al.*, 2016) Mediterranean waters.

**Supplementary Alignment S8**

Alignment of LBVG_174 and closely related genomes found in the EVG database, all of which were recovered from TARA Ocean Virome (Brum *et al.*, 2015).

**Supplementary Alignment S9**

Alignment of the cyanovirus S-EIVl (KJ410740) and its closely related genomes assembled from pelagic Arabian Sea waters (TARA_ERS488929_N000037) and from Lake Biwa (LBVG_9, _10, and _12).

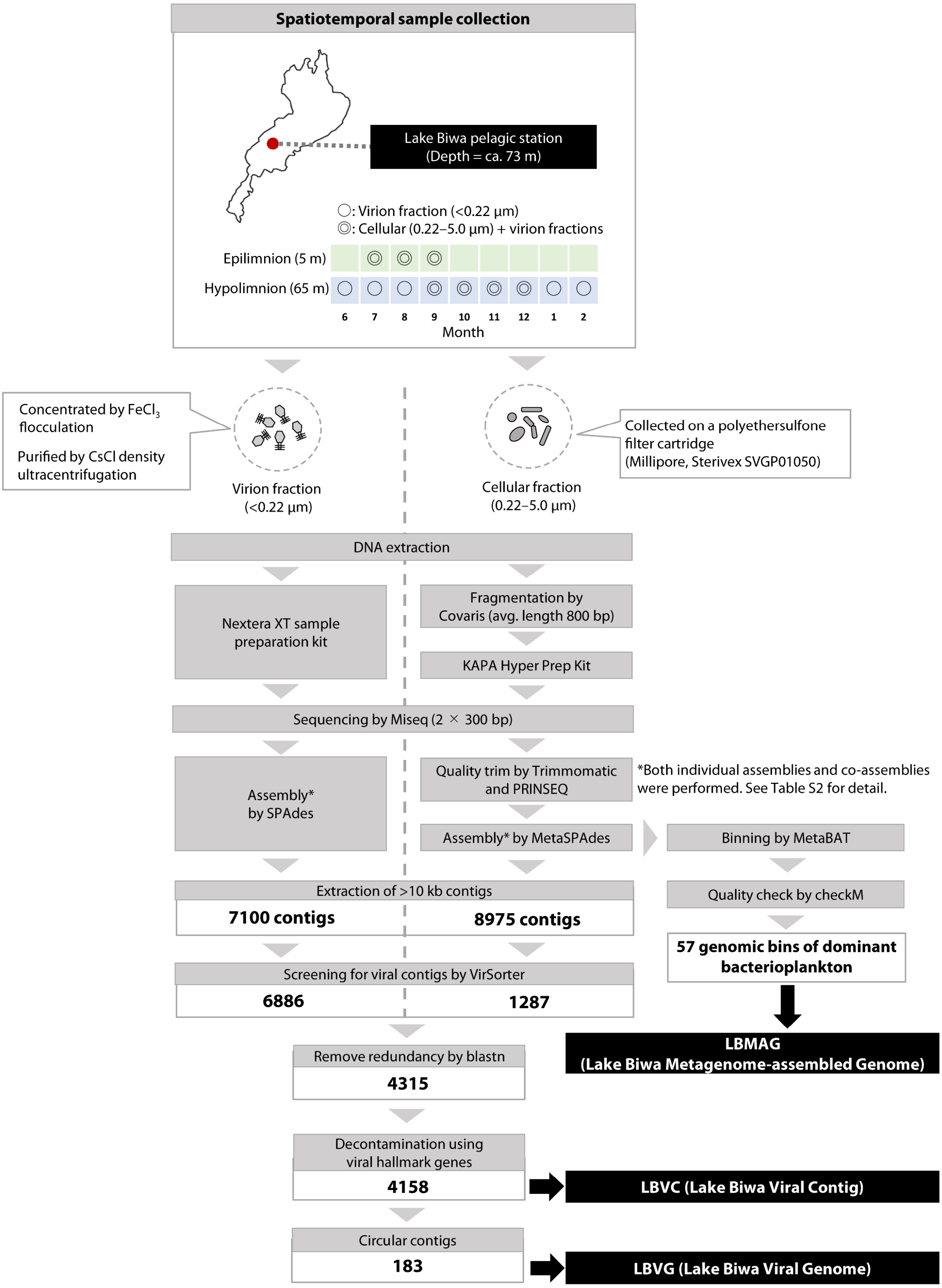


***Figure S1.*** Workflow of metagenomic sampling, sequencing, assembly, and creation of the Lake Biwa genomic datasets.

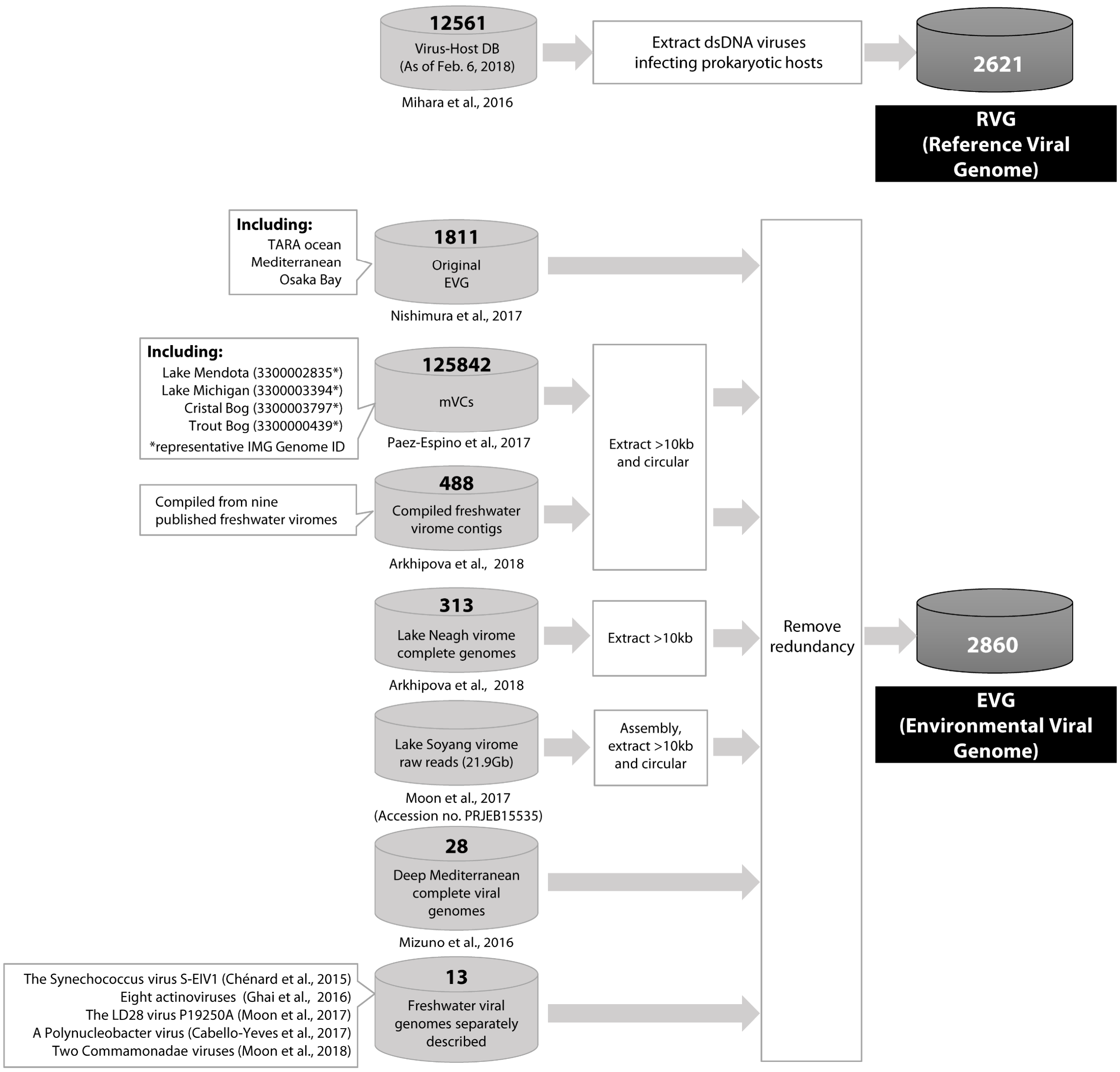


***Figure S2.*** Workflow for creation of the databases used in the study: reference viral genomes (RVGs) and environmental viral genomes (EVGs).

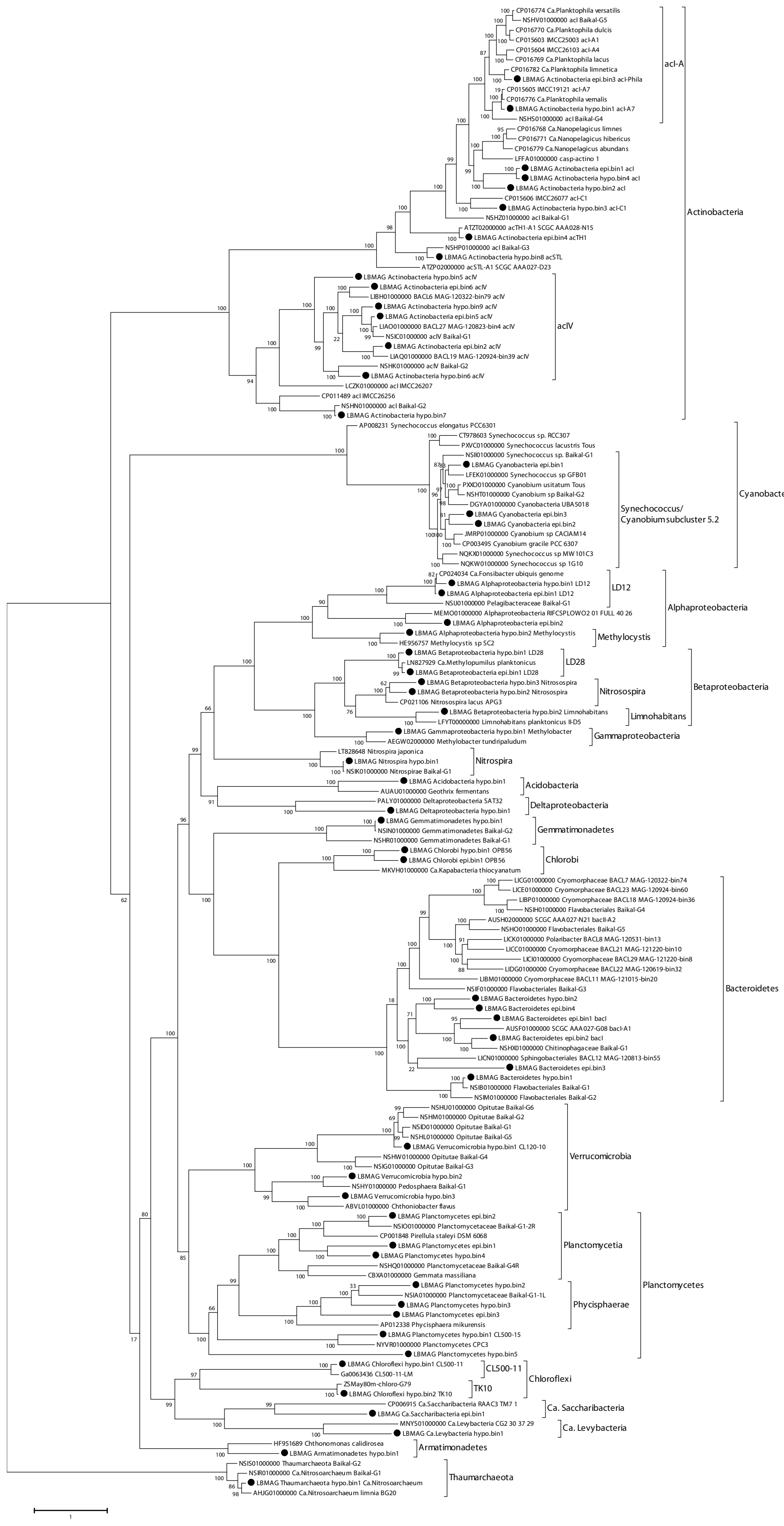


***Figure S3.*** A genome-resolved phylogenetic tree of Lake Biwa Metagenome Assembled Genomes (LBMAGs, black circles) and closely related published genomes, mainly derived from freshwater bacterioplankton. The tree was drawn based on conserved single-copy genes selected using the PhyloPhlAn software.

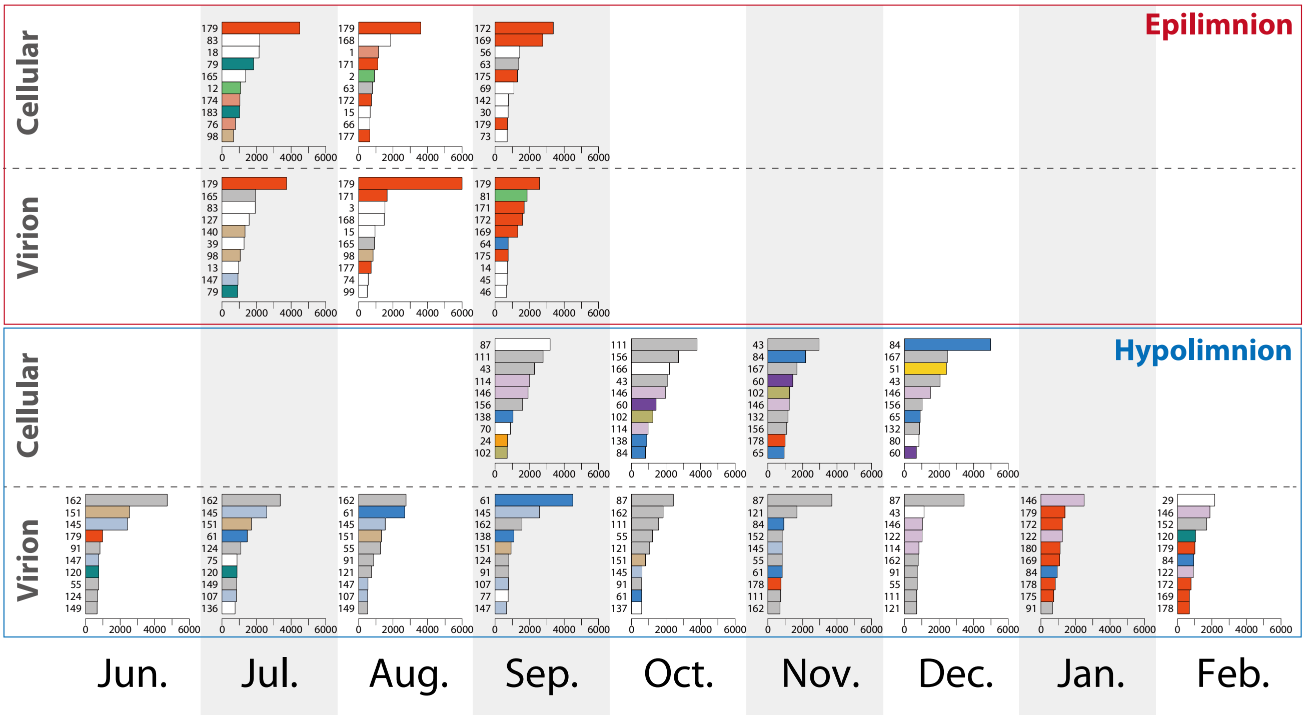


***Figure S4.*** Relative abundances (FPKM) of the 10 most predominant LBVGs in each sample. LBVG order and bar colors correspond to those shown in Figure 5.

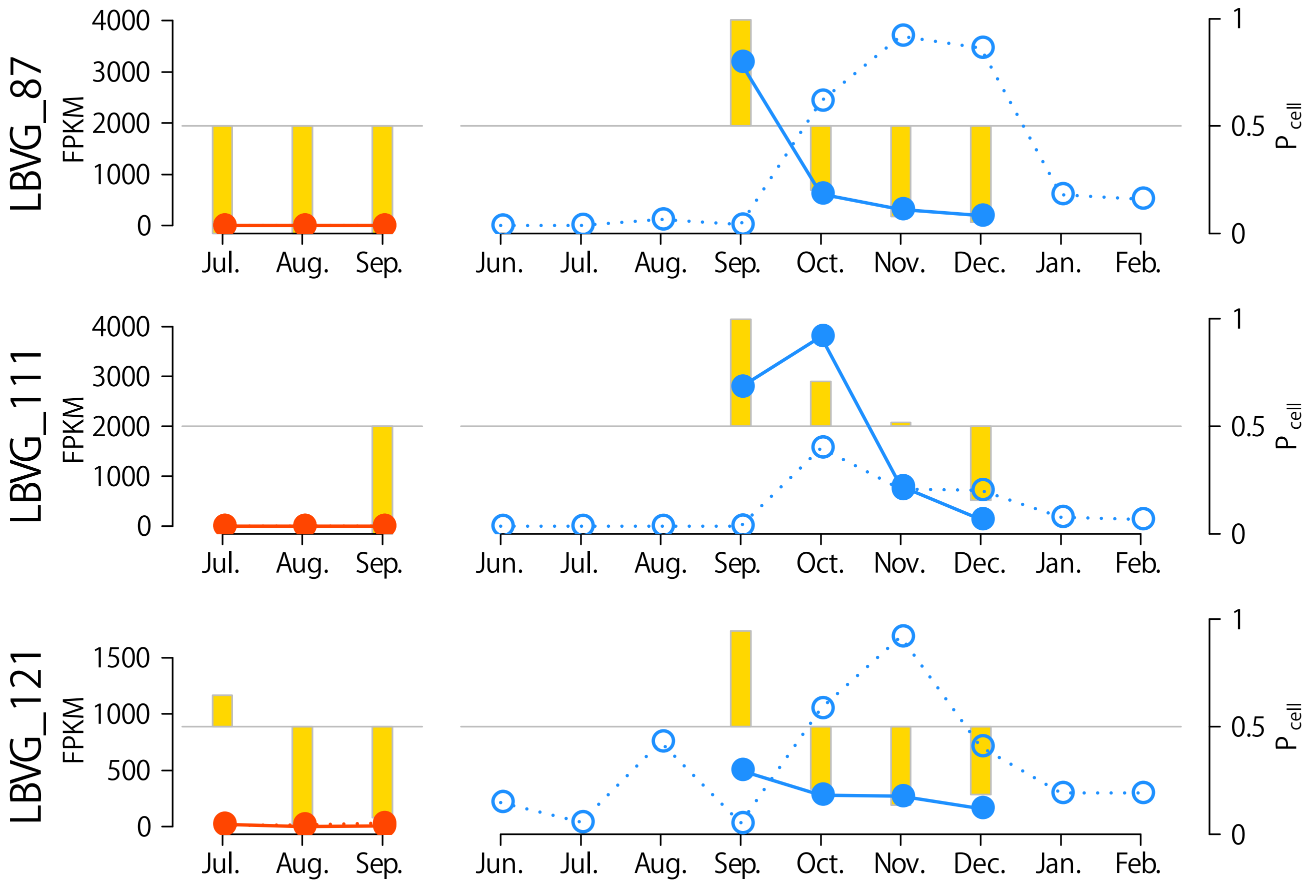


***Figure S5.*** Abundance succession of LBVG_87, _111, and _121, revealing their lysogenic induction in the hypolimnion. Left panels (red plots) and right panels (blue plots) indicate succession in the epilimnion and hypolimnion, respectively. Abundances within the cellular and virion fractions are indicated by solid and open points, respectively. Yellow bars indicate cellular–virion preference ($P_{\mathrm{cell}}$).

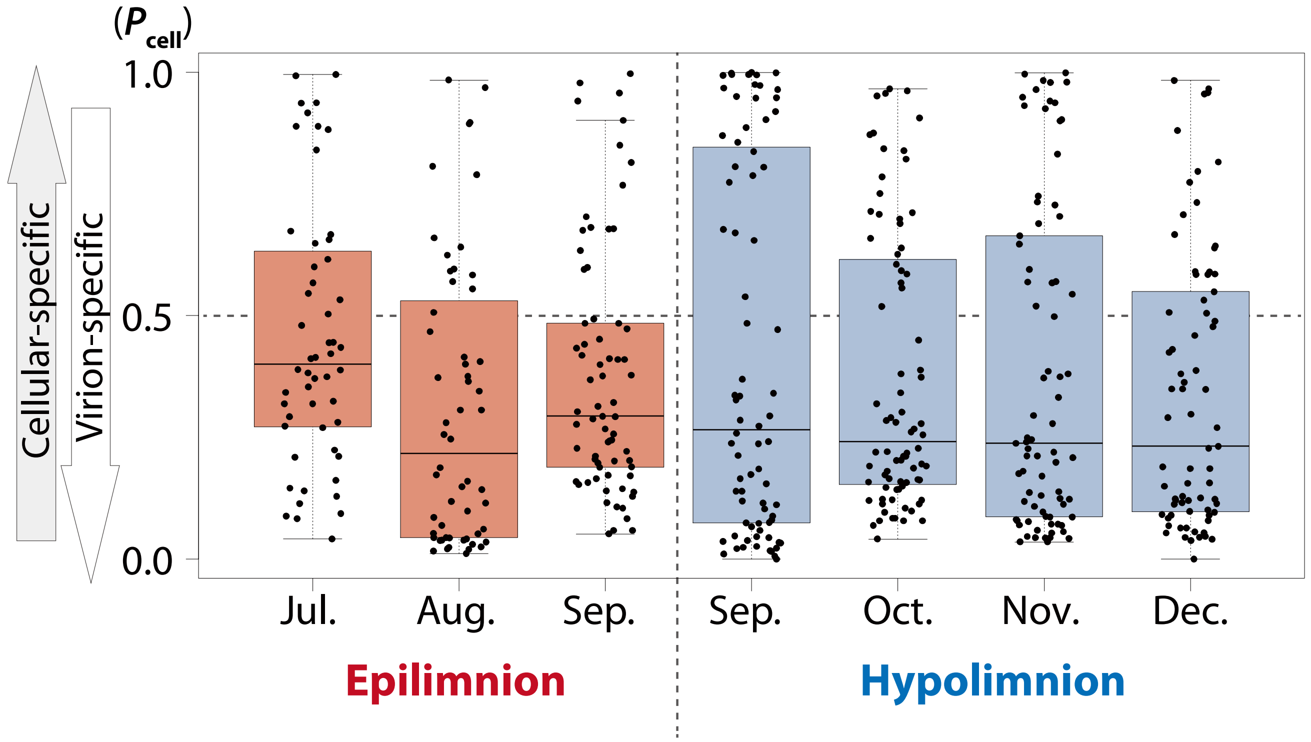


Pairs showing p<0.05 by the Wilcoxon rank sum test

| | | | |
|---|---|---|---|
| Aug. Epilimnion | < | Jul. Epilimnion | p = 0.002 |
| Aug. Epilimnion | < | Sep. Epilimnion | p = 0.021 |
| Oct. Hypolimnion | < | Jul. Epilimnion | p = 0.032 |
| Nov. Hypolimnion | < | Jul. Epilimnion | p = 0.041 |
| Dec. Hypolimnion | < | Jul. Epilimnion | p = 0.009 |
| Aug. Epilimnion | < | Oct. Hypolimnion | p = 0.038 |

***Figure S6.*** Comparison of cellular–virion preference ($P_{cell}$) among samples. Each point indicates an individual LBVG; its distribution is shown by a boxplot. Because $P_{cell}$ fluctuates greatly when abundance is low, LBVGs for which the sum of cellular and virion FPKM is <100 were omitted from this analysis.

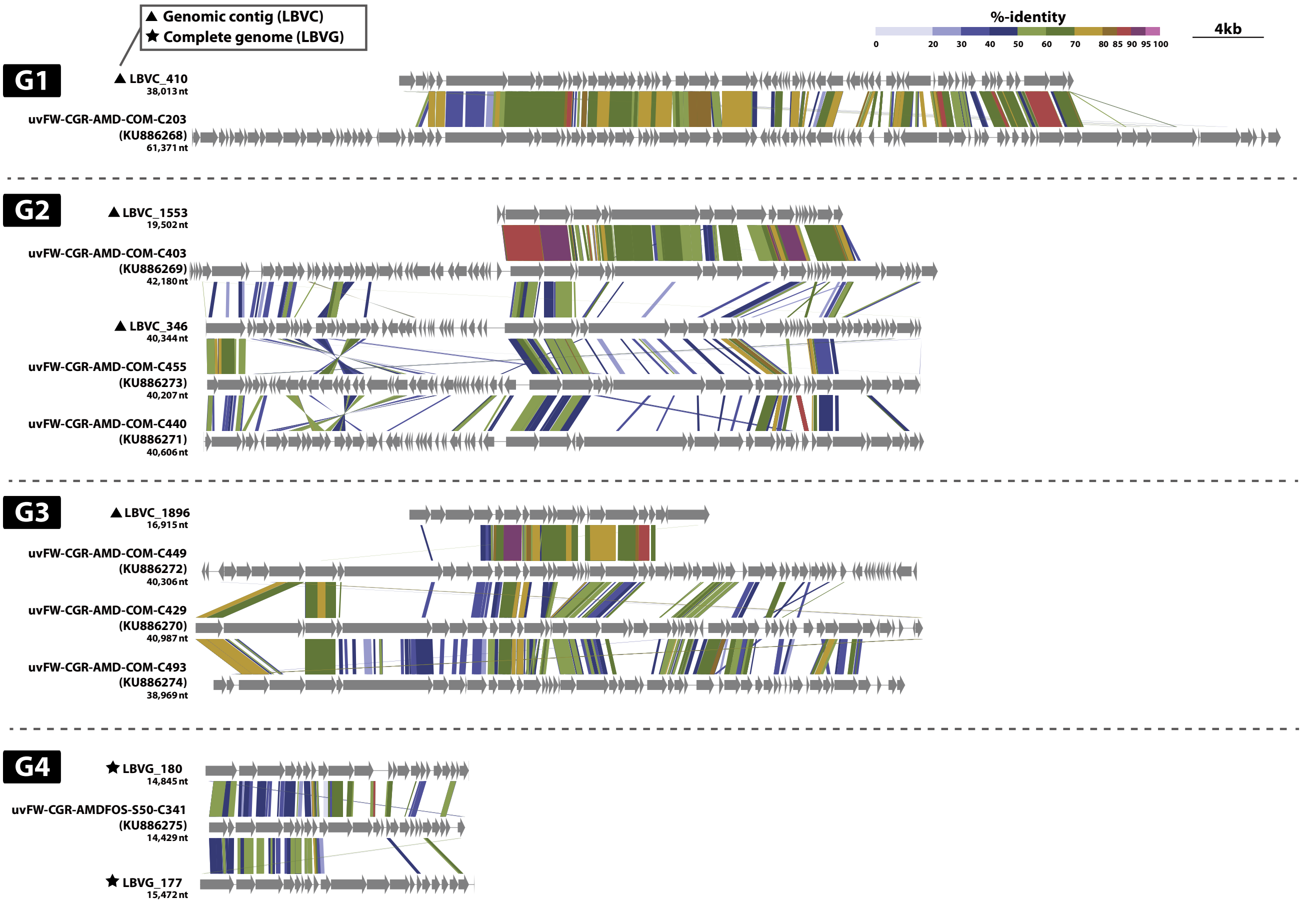


***Figure S7.*** tBLASTx alignments of the members of the four actinoviral groups described by Ghai *et al.* (2017) and their closely related sequences assembled in the present study, including LBVGs and LBVCs. Note that each genome was reversed and/or circularly permutated to improve alignment visibility.

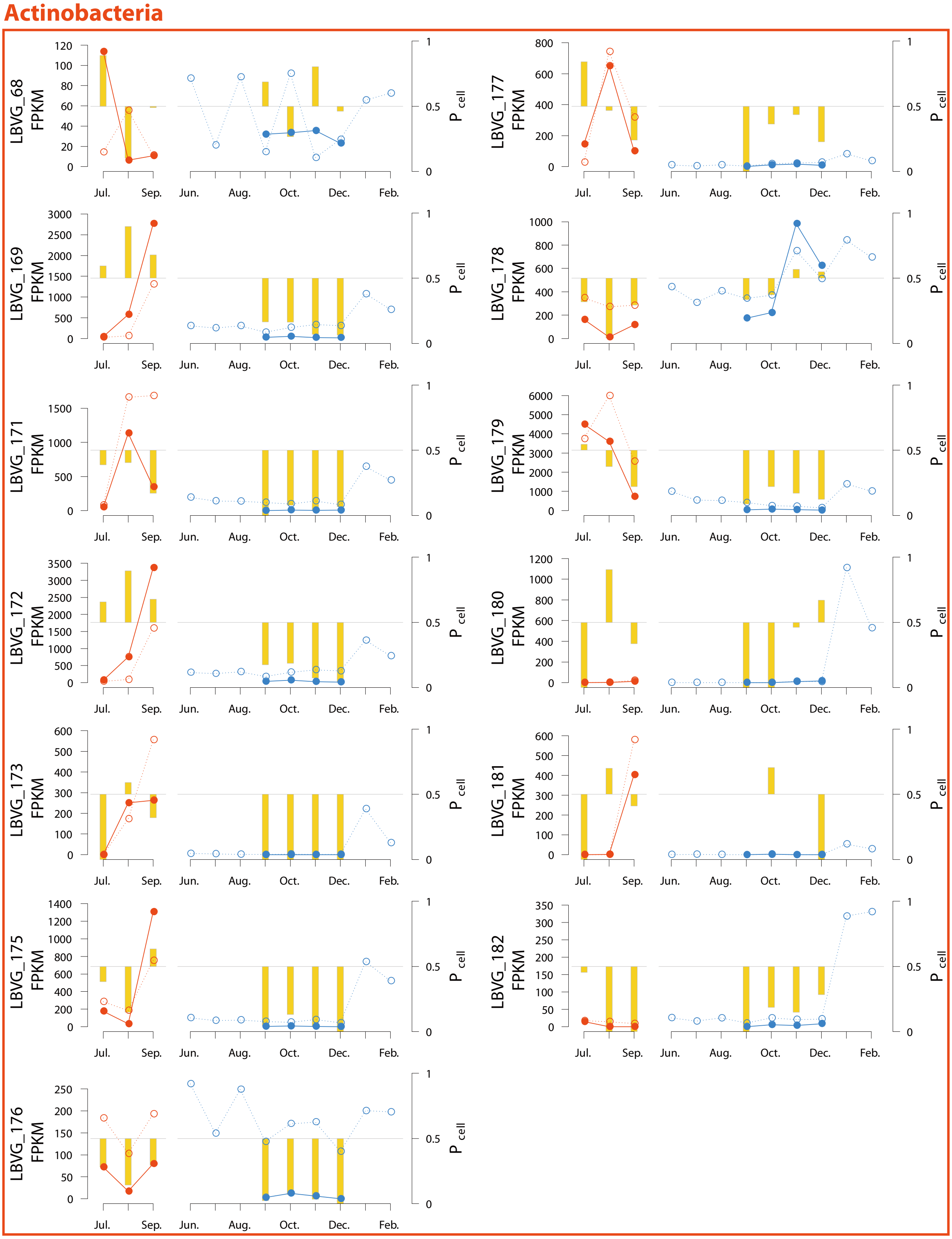


***Figure S8.*** Abundance succession of the 40 LBVGs with host predictions. LBVGs were sorted and grouped by the taxonomic affiliation of the host. Left panels (red plots) and right panels (blue plots) indicate succession in the epilimnion and hypolimnion, respectively. Abundances within the cellular and virion fractions are indicated by solid and open points, respectively. Yellow bars indicate cellular–virion preference ($P_{\mathrm{cell}}$).

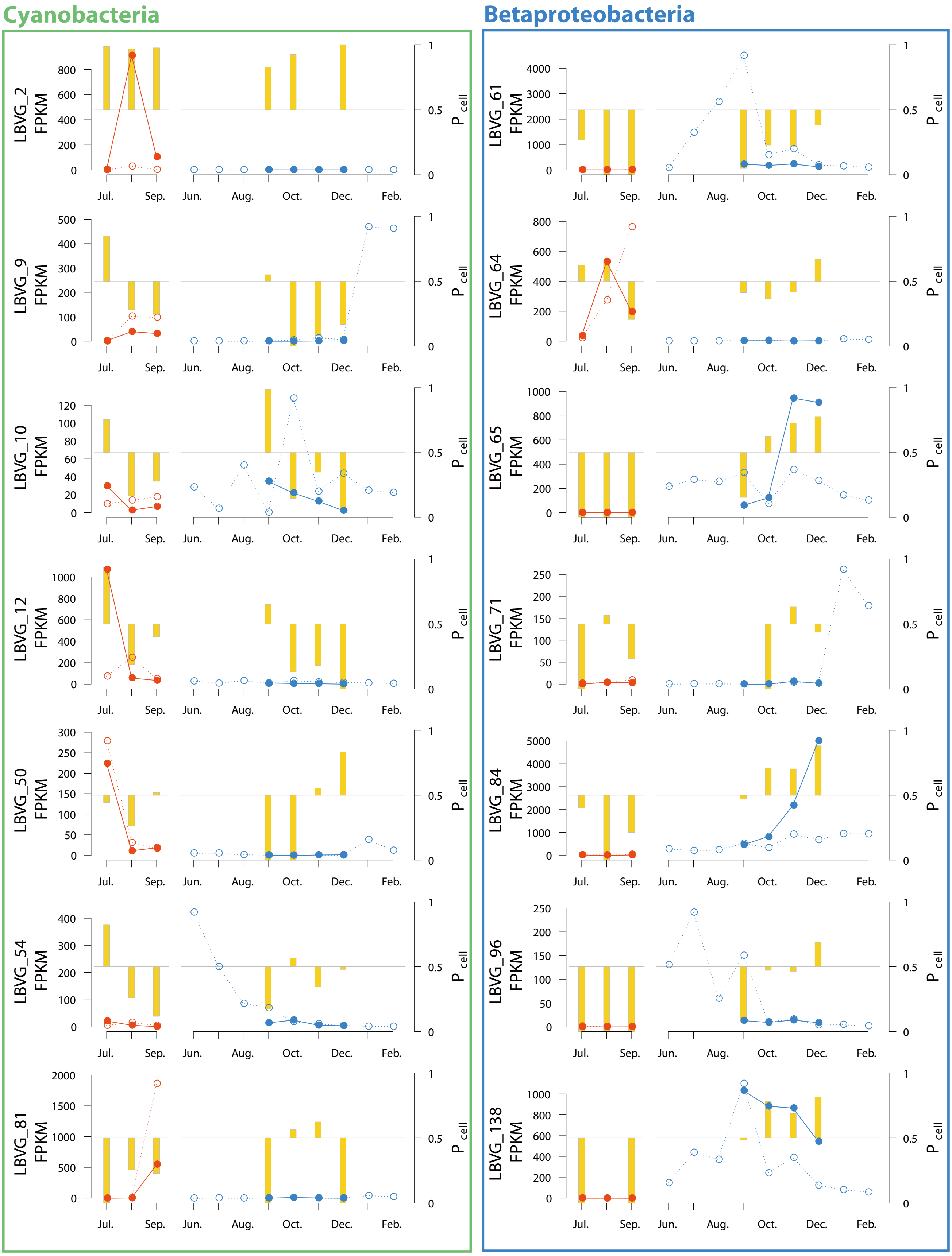


***Figure S8.*** Continued.

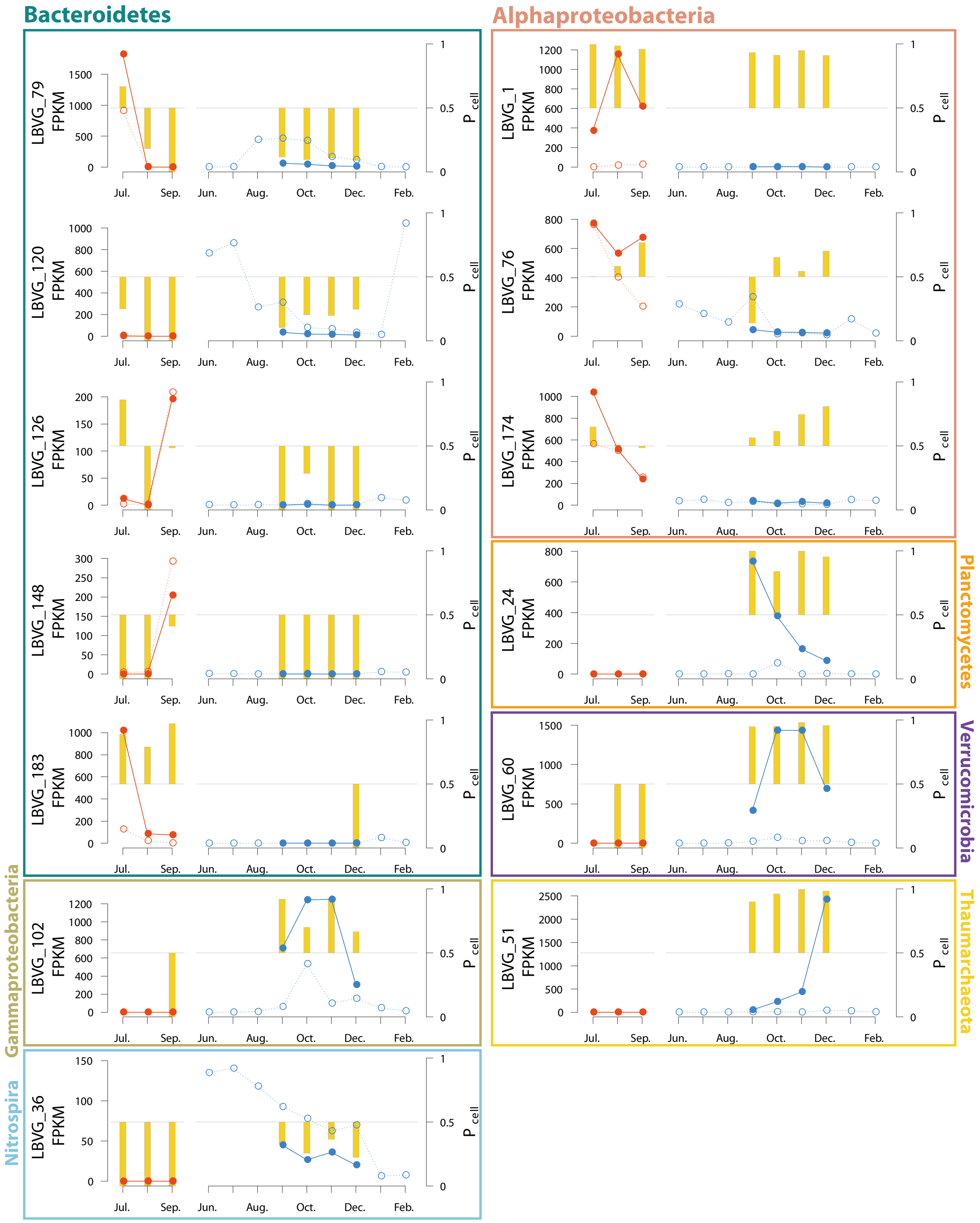


***Figure S8.*** Continued.

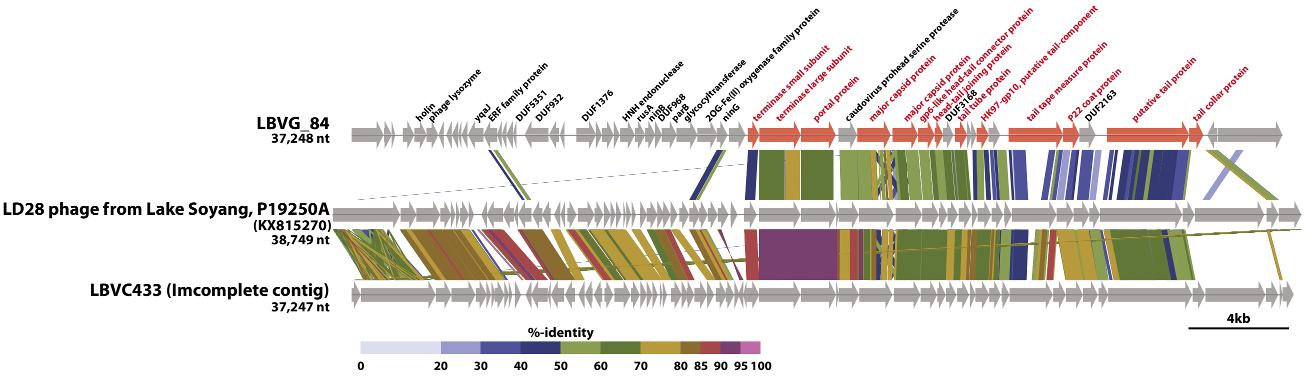


***Figure S9.*** tBLASTx alignment of the LD28 virus P19250A and its closely related sequences assembled in the present study, LBVG_84 (circular genome) and LBVC_433 (non-circular genomic contig). Functionally annotated genes in LBVG_84 are labeled. Annotations are red for viral structural genes, which were preferentially conserved among the genomes. Note that each genome was reversed and/or circularly permutated to improve alignment visibility.

### Captions for other Supplementary Information Files:

**Supplementary Table S1**

Environmental profile and number of sequencing reads generated for the 12 samples collected in the present study.

**Supplementary Table S2**

Sources and outputs of the assemblies in the present study. Each row represents one assembly; black circles indicate the source(s) of each (co-)assembly. Assemblies used to reconstruct LBVCs/LBVGs and LBMAGs are indicated in the contig source columns.

**Supplementary Table S3**

LBMAG profiles. Completeness and contamination were evaluated using the checkM software. Estimated genome size was calculated based on assembly size and completeness. Note that taxonomic assignment by GTDB-tk was based on the latest GTDB database at the time of writing (release 86) and may contain inconsistencies with the conventional taxonomic scheme (e.g., members of Betaproteobacteria were assigned to class Gammaproteobacteria, and *Ca*. Nitrosoarchaeum was assigned to phylum Crenarchaeota, class Nitrososphaeria).

**Supplementary Table S4**

Relative abundance (as FPKM) of individual LBMAGs in each sample.

**Supplementary Table S5**

Summary of evidence for host predictions. See the Workflow for host prediction section in Supplementary Materials and Methods for details.

**Supplementary Table S6**

A distance matrix of 6-mer nucleotide composition between the 183 LBVGs and 57 LBMAGs. Distance is indicated by the dissimilarity measure $d^*_2$, which was calculated by VirHostMatcher v. 1.0.0 (Ahlgren *et al.*, 2017). Values <0.25 are emphasized using white letters.

**Supplementary Dataset S1**

A summarized dataset generated in the present study. The Excel file consists of four worksheets: (i) LBVG: compiled data associated with 183 individual LBVGs, including genome size, GC content, habitat preference (as shown in Figure 1), gOTU affiliation, close relatives detected in the RVG and EVG databases, origin of close relatives in the EVG databases, relative abundance (as FPKM), presence of viral hallmark genes, and selected AMGs mentioned in the main text. (ii) LBVC: dataset for 4,158 individual LBVCs. Note that an $S_G$ value calculated for an incomplete partial genomic contig does not represent whole-genome similarity, and therefore is not comparable with values calculated between complete genomes. (iii) LBVG_ORFs: compiled data associated with 11,246 individual ORFs predicted in the 183 LBVGs, including functional annotations by DIAMOND to UniRef90, by hmmscan to pVOG, by eggnog-Mapper, and by pipeline_for_high_sensitive_domain_search. (iv) LBVC_ORFs: dataset for 127,961 individual ORFs predicted among the 4,158 LBVCs. The original nucleotide and amino acid sequences of LBVGs and LBVCs are available from https://doi.org/10.6084/m9.figshare.7934924.
